# Transient residence of the repulsive client Shutdown in Yb bodies plays a critical role in Piwi-piRISC biogenesis and maintaining fertility

**DOI:** 10.1101/2024.11.13.623522

**Authors:** Shigeki Hirakata, Aoi Fujita, Mikiko C. Siomi

**Affiliations:** Department of Biological Sciences, Graduate School of Science, The University of Tokyo, Tokyo 113-0032, Japan

**Keywords:** Shutdown, Yb bodies, Piwi-piRISC, piRNA, Armitage, liquid-liquid phase separation

## Abstract

Nonmembranous organelles, such as Yb bodies in *Drosophila* ovarian somatic cells, form through liquid-liquid phase separation of scaffold proteins. Client proteins transiently accumulate in these organelles by binding to scaffolds. However, the regulation of this process is not well understood. Here, we investigate Shutdown, a client protein in Yb bodies, the site of Piwi-piRISC precursor (pre-Piwi-piRISC) formation. We found that Shutdown connects Armitage and Piwi in the cytosol to promote Piwi deposition to Yb bodies, before quickly returning to the cytosol. This return allows Armitage to transfer pre-Piwi-piRISC to the mitochondria for maturation. Shutdown’s self-repulsion via its acidic N-terminus enables its transient residence in Yb bodies. A point mutation in Armitage, similar to a human Armitage/MOV10L1 mutation associated with azoospermia, traps Shutdown in Yb bodies, blocking Piwi-piRISC generation. This study reveals the mechanism of Shutdown’s transient localization in Yb bodies and its essential role in Piwi-piRISC production and maintaining fertility.

## Introduction

Intracellular liquid-liquid phase separation (LLPS) has emerged as a crucial mechanism for cellular organization and the facilitation of biochemical processes. LLPS leads to the formation of nonmembranous organelles, often referred to as biomolecular condensates or droplets, which play an integral role in various cellular functions^1–4^. These condensates are typically composed of two types of proteins: scaffold proteins, which form the structural basis of the droplet, and client proteins, which are recruited to the droplets in a manner dependent on the scaffold proteins^5^.

The formation and maintenance of biomolecular condensates are governed by multivalent interactions among scaffold proteins and, in some instances, nucleic acids. These interactions can be mediated through intrinsically disordered sequences or globular domains of the proteins^1–4^. The liquid nature of biomolecular condensates makes them ideal for the rapid entry and exit of materials, including client proteins^6^.

While the mechanisms governing client protein entry into droplets have been extensively studied^5,7–10^, those governing the exit of client components remain less understood. Recent studies have elucidated certain aspects of this process, including the involvement of HYL1 proteins in the release of mature miRNA from plant D-bodies^11^ and the RNA-binding-dependent release of Piwi-piRISC precursors (pre-Piwi-piRISC) from *Drosophila* Yb bodies^12^. From a more physical perspective, mature rRNA, the final product of rRNA assembly in the nucleolus, has been shown to be thermodynamically excluded from the nucleoli through a reduction in rRNA valency^13^. Notably, these reports indicate that products of RNA and RNA–protein (RNP) maturation processes, such as miRNAs, pre-Piwi-piRISC, and rRNAs, are released from nonmembranous organelles after processing, leading to a reduction in the concentration of these products within the droplet. In theory, this type of molecular dynamics promotes reactions by shifting the molecular equilibrium back to initial conditions. However, the effects of such product release from condensates on the physiological functions of these droplets have not yet been experimentally validated. Moreover, mechanisms governing the exclusion of client components from the condensates remain to be clarified.

In the context of the piRNA pathway, which plays a critical role in maintaining genomic integrity in reproductive tissues^14–19^, Yb bodies represent a notable example of biomolecular condensates within *Drosophila* ovarian somatic cells (OSCs)^20–24^. Yb bodies are formed via LLPS of the RNA-binding protein Female sterile (1) Yb (Yb) and *flam* RNA, the major source of piRNAs in OSCs^8^. These cytosolic droplets act as assembly sites for pre-Piwi-piRISC, composed of Piwi protein and *flam* in OSCs^12^, playing a vital role in the production of piRISC that targets transposons for silencing^8^. Indeed, the absence of the scaffold protein Yb results in the derepression of transposons, leading to infertility^20,22,25^.

Yb recruits Armitage (Armi), an RNA helicase, to Yb bodies, allowing Armi to bind to *flam* transcripts^20,22,23,26^. In addition, empty Piwi is recruited to Yb bodies in a manner dependent on Armi^12^, but the detailed molecular mechanisms remain unclear. *flam* transcripts undergo initial digestion by unknown nuclease(s), generating multiple 5′ ends to which Piwi binds, producing multiple pre-Piwi-piRISC. These molecular interactions lead to the formation of Armi-pre-Piwi-piRISC complexes, which exit Yb bodies and migrate to the outer mitochondrial membrane for Piwi-piRISC maturation by the endonuclease Zucchini (Zuc), located on the mitochondrial surface^12,20,26–29^.

Other known client proteins in Yb bodies, besides Armi and Piwi, include Sister of Yb (SoYb), Vreteno (Vret), and Shutdown (Shu)^30,31^. These proteins are essential for transposon repression in the ovaries^30–33^, although their specific molecular roles in the piRNA pathway are not yet fully understood. While Vret and SoYb form a heterodimer and accumulate in Yb bodies, Shu appears to be a transient component as it is largely dispersed in the cytosol in most OSCs^8^, although it is predominantly detected in Yb bodies in a subset of OSCs^31^.

Shu possesses a peptidyl-prolyl isomerase (PPIase) domain with mutations in the active site, rendering it enzymatically inactive but potentially functioning as a protein-protein interaction domain^31,33,34^, as observed in other proteins under similar circumstances^35–37^. However, its partner proteins have yet been identified. Shu also has a tetratricopeptide repeat (TPR) domain, known to bind to heat shock proteins such as HSP90 in mammals (HSP83 in *Drosophila*)^38,39^. While Shu in the germline was reported to cooperate with HSP83^31^, its physical binding to HSP83 remains an open question. By examining the interactions between Shu and other components of Yb bodies, we can uncover the intricate balance of attractive and repulsive forces that govern protein localization to Yb bodies. This provides a model for investigating the dynamics of transient clients within cellular droplets and allows the elucidation of its physiological significance.

In this study, we first focused on identifying the binding partners of Shu in cultured OSCs. We found that Shu does not bind to HSP83 but interconnects Armi and cytosolic Piwi, facilitating Piwi transport to Yb bodies along with Armi. Inside the bodies, the SoYb/Vret heterodimer receives Piwi from Shu, supporting the assembly of pre-Piwi-piRISC. Thus, in the absence of Shu, Piwi fails to become pre-piRISC and remains wandering in the cytosol. We also found that the N-terminus of Shu has an acidic region that imparts a repulsive property, preventing it from self-associating. Exploiting this property, Shu returns to the cytosol immediately after Piwi deposition in Yb bodies. This allows the exit of pre-Piwi-piRISC from Yb bodies, facilitating Zuc-mediated Piwi-piRISC maturation. A recent human genome study discovered the S816I point mutation in MOV10L1, the mouse counterpart of Armi, in a patient with nonobstructive azoospermia. The present study shows that the *Drosophila* Armi S769I mutant, which corresponds to the MOV10L1 S816I mutant, causes Armi to adhere to Shu, retaining Shu and Piwi in Yb bodies and thereby inhibiting Piwi-piRISC generation. The biological significance of the release of transient client Shu from Yb bodies, mediated by its repulsive N-terminus, and its direct association with infertility is now clear.

## Results

### Shu delivers Piwi into Yb bodies through interaction with Armi

To clarify the binding of Shu with other clients in Yb bodies, we first generated a polyclonal antibody against Shu (Figure S1A). Using this antibody, the Shu complex was immunoprecipitated from cultured OSCs, and the presence of Piwi, Armi, SoYb, and Vret in the complex was examined by western blotting; Piwi, but not Armi and the SoYb/Vret heterodimer, was detected in the Shu complex (Figure 1A). When OSC lysates lacking Shu were alternatively used, Piwi was barely detected (Figures 1A and S1B), verifying the Shu–Piwi interaction. The association between Shu and Piwi was maintained in the absence of Armi, SoYb, and Vret (Figure 1B), indicating that Shu binds to Piwi in a manner independent of Armi and the SoYb/Vret heterodimer. No HSP83 signal was detected in the Shu complex (Figure S1C), suggesting that Shu may not bind to HSP83 in OSCs, although Shu has a TPR domain, which is known to bind heat shock proteins. In our previous study, Piwi was detected in the Armi complex^20^, which was confirmed in the current study (Figure S1D). This interaction occurred independently of Yb, indicating that Piwi and Armi can interact outside of Yb bodies, whose assembly depends on the scaffold Yb. The Piwi complex also contained Armi (Figure 1C), but in the absence of Shu, Armi was no longer associated with Piwi (Figure 1C). These results suggest that the Piwi–Armi association depends on Shu, although the Piwi–Shu interaction is independent of Armi.

**Figure 1.**
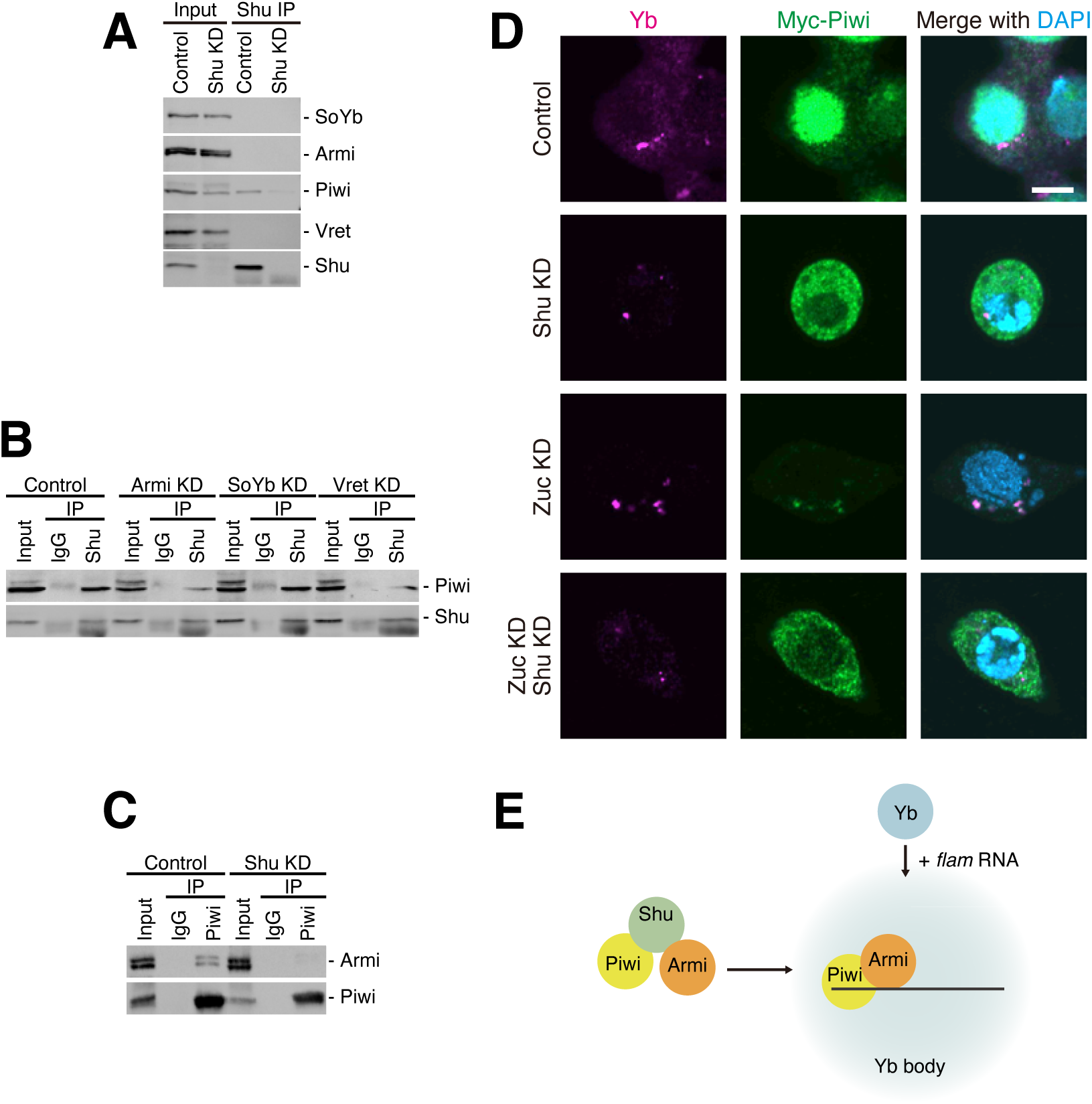
Shu transports Piwi to Yb bodies. (A) The Shu complex was immunoisolated from OSCs using anti-Shu antibodies and was found to contain Piwi. Shu-depleted OSCs (Shu KD) served as a negative control. (B) Shu and Piwi remained bound to each other upon depletion of Armi, SoYb, or Vret. Non-immune IgG served as a negative control. (C) Upon Shu depletion, Piwi no longer associated with Armi. Non-immune IgG served as a negative control. (D) Immunofluorescence analyses in OSCs. Upon Shu depletion, Myc-tagged Piwi (Myc-Piwi) failed to enter the nuclei and dispersed in the cytosol. Myc-Piwi accumulated in Yb bodies upon Zuc depletion^20,22^; however, with co-depletion of Zuc and Shu, Myc-Piwi dispersed in the cytosol instead. Nuclei are shown in blue. Scale bar: 5 μm. (E) Summary of the molecular function of Shu in OSCs. Yb protein and *flam* RNA form Yb bodes via LLPS. Shu recruits Piwi to the granule through interaction with Armi. See also Figure S1.

A previous study showed the interaction of Shu with Armi and Piwi in OSCs^31^. We performed immunoprecipitation using FLAG-tagged Shu (Shu-FLAG) expressed in OSCs and found that Armi co-immunoprecipitated with the exogenous Shu (Figure S1E), confirming the Armi–Shu interaction. The Armi–Shu interaction was found to be independent of Piwi and the SoYb/Vret heterodimer (Figures S1F and S1G).

Piwi is localized to the nucleus upon its association with piRNAs, but unloaded Piwi remains to be cytosolic^40,41^. In normal OSCs, Myc-tagged Piwi (Myc-Piwi) was detected in the nucleus, indicating that Myc-Piwi is bound with piRNAs and is capable of transposon silencing in the nucleus, similarly to endogenous Piwi (Figure 1D). However, upon Shu depletion, Myc-Piwi became almost exclusively cytosolic, although the expression level of the exogenous Piwi was fairly maintained (Figures 1D and S1H). This validates the importance of Shu in Piwi-piRISC generation and subsequent transposon silencing.

Piwi becomes pre-Piwi-piRISC in Yb bodies and then migrates with Armi to the mitochondrial surface, where the endoribonuclease Zuc cleaves the Piwi-bound piRNA precursor, liberating mature Piwi-piRISC^12,27–29^. When piRISC maturation was inhibited by the loss of Zuc, Piwi accumulated in Yb bodies^20,22^ (Figure 1D). We next depleted Shu under these conditions and found that Piwi signals were almost exclusively scattered in the cytosol (Figure 1D). This suggests that Shu is responsible for concentrating cytosolic Piwi in Yb bodies. We previously reported that Armi is also necessary for transferring Piwi to Yb bodies, as the loss of Armi rendered Piwi cytosolic^20,22^. However, the roles of Shu and Armi at this step seem to differ from each other; Shu interconnects Armi and Piwi, while Armi transfers the Piwi–Shu complex to Yb bodies (Figure 1E). The subcellular localization of Armi was minimally affected by Shu depletion (Figure S1I), supporting this notion.

### Shu deposits Piwi into Yb bodies and immediately returns to the cytosol

Previous studies have not reached a consensus regarding the subcellular localization of exogenous Shu in OSCs^8,31,33^. In this study, we immunostained naïve OSCs with anti-Shu antibody. Approximately 10% of the cells in the same field of view contained Shu-positive Yb bodies (Figures 2A and 2B). This confirms that Shu is a transient resident of Yb bodies and suggests a dynamic movement of Shu *in vivo*: Shu returns to the cytosol shortly after delivering its cargo in Yb bodies.

**Figure 2.**
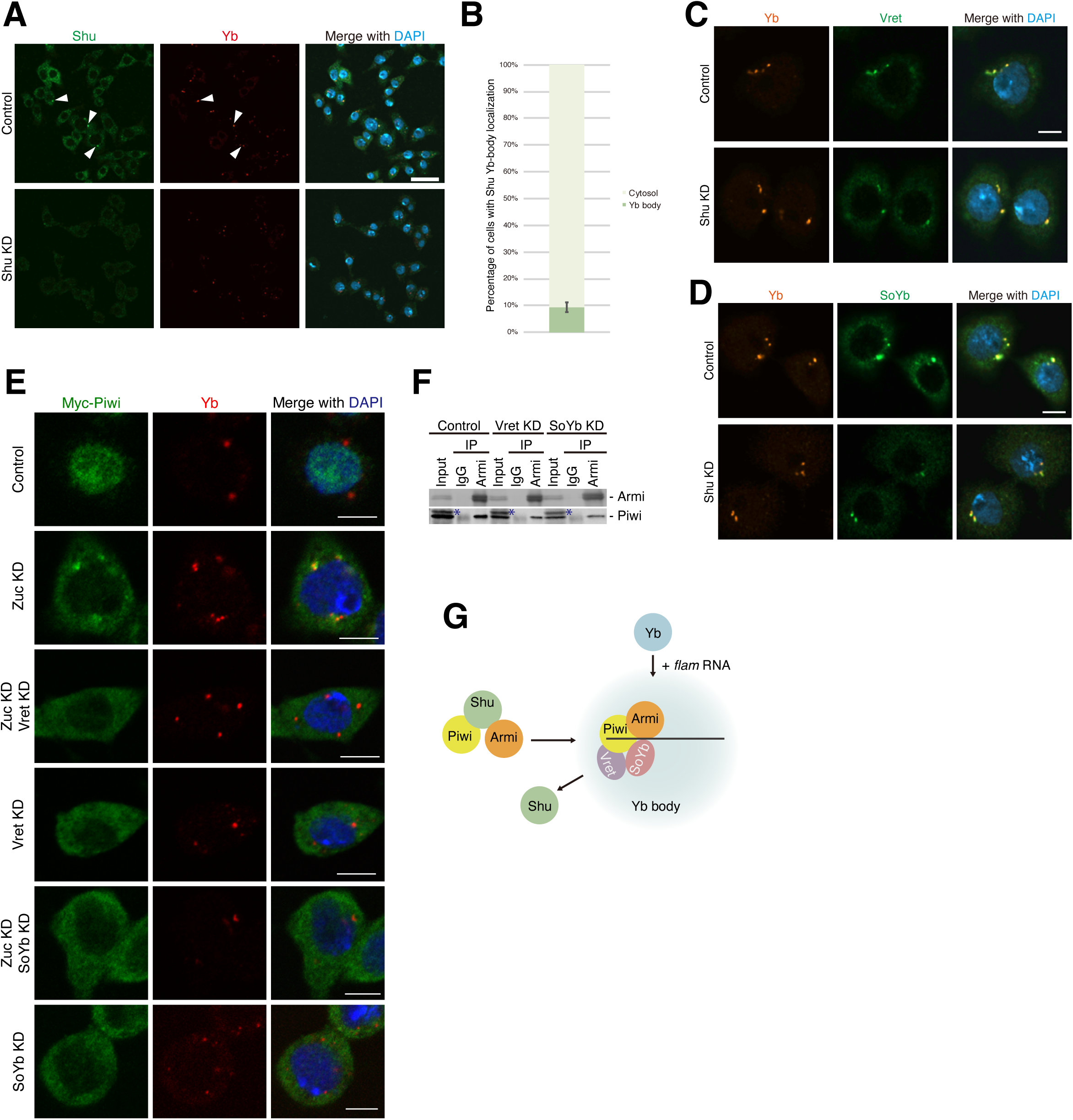
The SoYb/Vret heterodimer receives Piwi from Shu in Yb bodies. (A, B) Immunofluorescence analyses in OSCs using anti-Shu antibodies. In most cells, Shu did not accumulate in Yb bodies (A). Quantification revealed that Shu was dispersed in the cytosol in 90% of cells (B). Yb bodies are indicated by white arrowheads. Nuclei are shown in blue. Scale bar: 15 μm. Over 100 cells were analyzed in each biological triplicate (> 300 cells in total). (C, D) Vret (C) and SoYb (D) were still localized in Yb bodies upon depletion of Shu. Nuclei are shown in blue. Scale bar: 5 μm. (E) Immunofluorescence analyses in OSCs. Myc-Piwi accumulated in Yb bodies upon Zuc depletion^20,22^. Upon depletion of Vret or SoYb, Myc-Piwi no longer accumulated in Yb bodies and dispersed in the cytosol, even in Zuc-depleted cells. Nuclei are shown in blue. Scale bar: 5 μm. (F) Depletion of Vret and SoYb had minimal effect on the interaction between Armi and Piwi. Non-immune IgG served as a negative control. Asterisks indicate nonspecific bands caused by the secondary antibody. (G) Model of protein entry and exit from Yb bodies. Piwi interacts with Armi in a Shu-dependent manner and is recruited to Yb bodies. However, Shu rarely accumulates in Yb bodies and returns to the cytosol. Vret and SoYb receive Piwi in Yb bodies, where pre-Piwi-piRISC is assembled. See also Figure S2.

Which factor(s) receive Piwi from Shu in Yb bodies? We speculate that the SoYb/Vret heterodimer is a feasible candidate, as it is a major, constant inhabitant of Yb bodies, though its function remains vague^8,30^. In fact, the subcellular localization of SoYb and Vret was not influenced by the loss of Shu and Piwi (Figures 2C, 2D, S2A, and S2B). As mentioned earlier, in the absence of Zuc, Myc-Piwi was abnormally concentrated in Yb bodies [Figures 1D, 2E (top six panels), and S2C]. To investigate the role of SoYb and Vret, we further depleted them under the circumstances: in both cases, Myc-Piwi was dispersed in the cytosol and was barely detected in Yb bodies (Figures 2E and S2C). The localization of Myc-Piwi was largely unaffected by Zuc restoration (Figures 2E and S2C). We previously reported that the Yb body localization of Armi is barely affected by depletion of SoYb and Vret^8^. Additionally, the Armi–Piwi interaction was maintained in the absence of SoYb and Vret (Figure 2F), indicating that the Shu-mediated supply of Piwi to Yb bodies persists under these conditions.

Based on these findings, we propose that Shu transports Piwi into Yb bodies, where the SoYb/Vret heterodimer receives Piwi from Shu, allowing Shu to return to the cytosol (Figure 2G). In this way, Shu can continuously transport Piwi to Yb bodies, facilitating the assembly of pre-Piwi-piRISC and the production of Piwi-piRISC.

### Repulsive property of the N-terminal acidic region of Shu is important for its return to the cytosol following Piwi delivery to Yb bodies

The N-terminal region of Shu, up to the PPIase domain, is rich in acidic residues, particularly between Asp41 and Glu73 (13 in 33 residues; 39.4%) (Figures 3A and S3A). Because amino acids of the same polarity, either acidic or basic, tend to repel each other when placed in close proximity, as in Yb bodies and other condensates, it was speculated that this repulsion drives Shu to leave Yb bodies after Piwi delivery.

**Figure 3.**
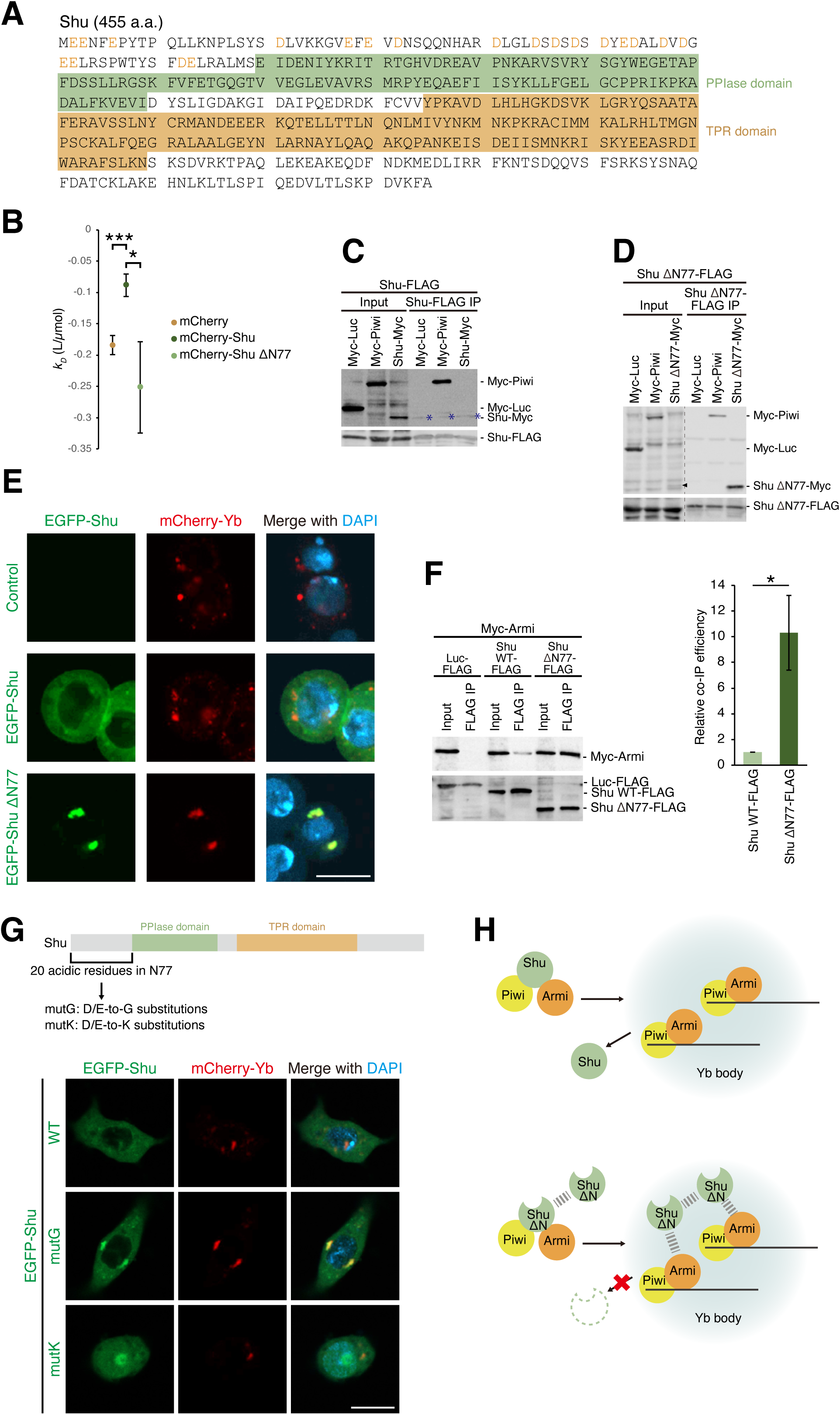
The N-terminal residues of Shu are repulsive and necessary for release of Shu from Yb bodies. (A) Amino acid sequence of Shu (CG4735). The PPIase and TPR domains are highlighted in green and orange boxes, respectively. Acidic residues in the N-terminal region are shown in orange. (B) Concentration dependence of diffusion coefficients (*k_D_*) of recombinant proteins. mCherry-tagged Shu (mCherry-Shu) exhibited repulsive behavior compared to mCherry alone and the N-terminal deletion mutant (ΔN77). Error bars represent standard errors calculated from 15 data points. *p < 0.05, ***p < 0.001. (C) FLAG-tagged Shu (Shu-FLAG) co-purified with Myc-Piwi but not with Myc-tagged Shu (Shu-Myc). Myc-tagged Luc (Myc-Luc) served as a negative control. Asterisks indicate IgG heavy chains. (D) Deletion of N77 allowed Shu-FLAG to co-purify with Shu-Myc in addition to Myc-Piwi. Myc-Luc was used as a negative control. (E) Deletion of N77 caused EGFP-tagged Shu (EGFP-Shu) to accumulate in Yb bodies. Nuclei are shown in blue. Scale bar: 10 μm. (F) The interaction between Shu-FLAG and Myc-tagged Armi (Myc-Armi) was stabilized upon deletion of N77 from Shu. The graph on the right shows quantified data. Error bars represent means ± SEM values from four independent experiments. *p < 0.05. FLAG-tagged Luc (Luc-FLAG) was used as a negative control. (G) Upper: twenty acidic residues in the N-terminal region were substituted with Gly (mutG) or Lys (mutK). Lower: mutG resulted in EGFP-Shu accumulation in Yb bodies, while mutK showed minimal accumulation in the granule, with some proteins localizing to the nuclei, likely due to the acquisition of a nuclear localization signal (NLS) through mutation. Nuclei are shown in blue. Scale bar: 10 μm. (H) Model of the physiological significance of Shu’s repulsive nature. Upper: wild-type Shu rapidly returns to the cytosol after depositing Piwi in Yb bodies. Lower: deletion of the N-terminus reduces Shu’s repulsion, leading to self-association and multivalent Armi binding (dashed lines) in Yb bodies, stabilizing interaction between Shu and Armi and inhibiting Shu’s dispersion. See also Figure S3.

We first confirmed the repulsive nature of Shu’s N-terminus through *in silico* predictions. Using FINCHES^42^, we predicted the regions of Shu engaged in repulsion or attraction between two Shu molecules, assuming these regions are disordered. This analysis indicated strong repulsion between the acidic N-terminus of two Shu molecules (Figure S3B). Notably, this region is indeed predicted to be disordered by metapredict V2-FF^43^ (Figure S3B), fulfilling the prerequisites for the FINCHES analysis. We then predicted the effect of N-terminal deletion on this repulsion. Given that the remainder of the Shu protein, excluding the N-terminus, is predicted to be non-disordered (Figure S3B), we employed FMAPB2 to assess changes in the potential of mean force (PMF) as a function of distances between two rigidly folded proteins^44^. The PMF of the deletion mutant ΔN77, which lacks Met1–Leu77 of Shu, but not wild-type (WT) protein, increased with greater interprotein distances, suggesting spontaneous attraction between two ΔN77 mutant proteins (Figure S3C).

To experimentally validate these predictions, we first measured the concentration dependence of diffusion coefficients (*k_D_*) of fluorescently labeled recombinant proteins. Proteins with lower *k_D_* values exhibit a more significant reduction in diffusion coefficients with increasing concentration, indicating stronger attractive forces between protein molecules. mCherry-tagged Shu (mCherry-Shu) displayed a higher *k_D_* value than both mCherry alone and the ΔN77 mutant (Figures 3B, S3D, and S3E), supporting the hypothesis that Shu’s N-terminal region confers repulsive properties.

Next, protein-protein interaction assays in OSCs showed that Shu has little to no ability to self-associate under conditions where the Shu–Piwi interaction was readily detected (Figure 3C). However, unlike Shu WT, Shu ΔN77 self-associated (Figures 3C and 3D). Subcellular localization analysis showed that EGFP-tagged Shu (EGFP-Shu) WT only slightly localized to Yb bodies, while EGFP-Shu ΔN77 accumulated significantly in Yb bodies (Figures 3E and S3F). These results support the hypothesis that the N-terminal acidity of Shu prevents its self-association and promotes its return to the cytosol after transporting Piwi to Yb bodies. Interestingly, the interaction between Shu and Armi was notably stabilized by the N-terminal deletion (Figure 3F). In the absence of the repulsive property, Shu’s self-association prolongs its residence in Yb bodies due to stabilized interactions with Armi, presumably driven by multivalent interactions induced by the self-association of Shu.

To further confirm this model, we generated two Shu substitution mutants: EGFP-Shu mutG, where all 20 acidic residues in the N-terminal region (Met1-Leu77) were substituted with glycine, and EGFP-Shu mutK, where all acidic residues were replaced by lysine (Figure 3G). The N-terminal repulsion between two mutG molecules was predicted to be much weaker than that of WT, while the repulsion between mutK molecules was predicted to be stronger than that of WT (Figures S3B and S3G). As expected, mutG, but not mutK, accumulated in Yb bodies (Figures 3G and S3H), further supporting the model that electrostatic repulsion between Shu molecules is essential for its release from Yb bodies (Figure 3H).

### Forced self-association of Shu impedes Piwi-piRISC generation by inhibiting the departure of pre-Piwi piRISC

To explore the impact of the loss of Shu’s repulsive nature on its function, we applied a nanobody strategy that leverages the tight interaction between EGFP and EGFP-binding protein (GBP)^45^. Shu was individually fused to both EGFP and GBP and expressed in OSCs (Shu-EGFP and Shu-GBP-FLAG) (Figure S4A). As references, GBP alone (GBP-FLAG) and EGFP alone (Myc-EGFP) were expressed when necessary (Figure S4A). We then examined the subcellular localization of these proteins in OSCs. When Shu-EGFP and Shu-GBP-FLAG were expressed individually, both fusion proteins accumulated only weakly in Yb bodies (Figure 4A). Coexpression of GBP-FLAG with Shu-EGFP or Myc-EGFP with Shu-GBP-FLAG hardly changed their localizations (Figure 4A). However, when Shu-EGFP and Shu-GBP-FLAG were coexpressed, both accumulated strongly in Yb bodies (Figure 4A). We argue that forced self-association of Shu causes it to reside much longer in Yb bodies. In the absence of Armi, this effect was eliminated (Figures 4B and S4B), supporting our earlier observation that Shu’s Yb body localization depends on Armi.

**Figure 4.**
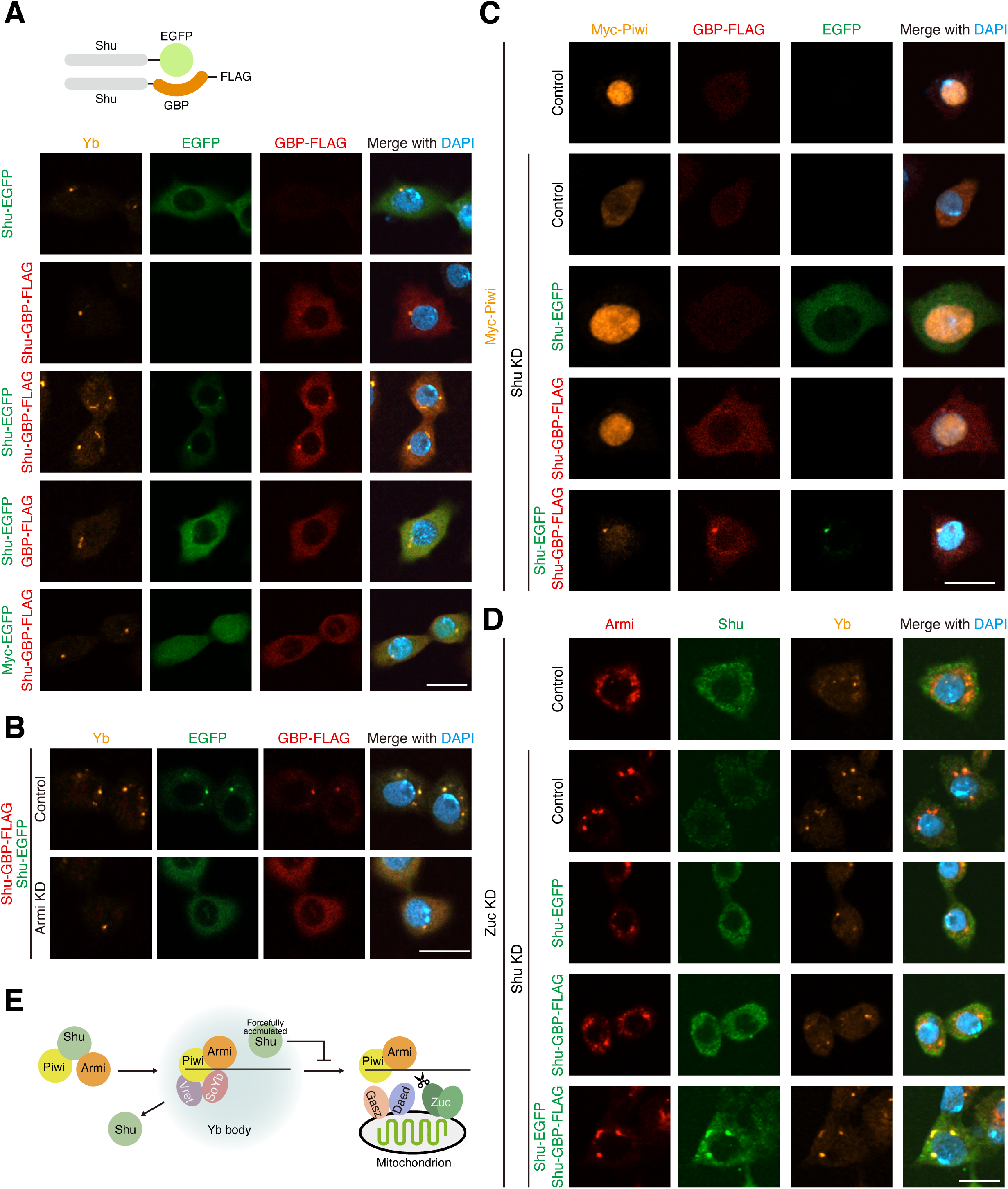
Dispersion of Shu from Yb bodies is required for pre-Piwi-piRISC departure toward mitochondria. (A) Upper: the affinity between Shu proteins was increased in OSCs using a nanobody strategy. Lower: coexpression of EGFP-tagged Shu (Shu-EGFP) and GBP-tagged Shu (Shu-GBP-FLAG) resulted in tagged-Shu accumulation in Yb bodies. GBP alone (GBP-FLAG) and EGFP alone (Myc-EGFP) were used as negative controls. Nuclei are shown in blue. Scale bar: 10 μm. (B) Accumulation of Shu-EGFP and Shu-GBP-FLAG in Yb bodies was lost upon depletion of Armi. Nuclei are shown in blue. Scale bar: 10 μm. (C) Rescue experiments showed that forced accumulation of Shu in Yb bodies hindered Piwi departure from Yb bodies. The accumulation of Piwi in Yb bodies, under condition where Shu’s return to the cytosol was blocked by forced accumulation, suggests that the return of Shu is not critical for Piwi recruitment, at least when Shu is over expressed. Nuclei are shown in blue. Scale bar: 10 μm. (D) Rescue experiments demonstrated that forced Shu accumulation in Yb bodies resulted in accumulation of Armi in Yb bodies in Zuc-depleted cells. Upon depletion of Zuc, Armi accumulates on mitochondria^12,29^. Depletion of Shu, which depletes Piwi from Yb bodies, caused Armi to accumulate in Yb bodies, even under the Zuc-depleted condition. Coexpression of Shu-EGFP and Shu-GBP-FLAG also led to Armi accumulation in Yb bodies in Zuc-depleted cells. Nuclei are shown in blue. Scale bar: 10 μm. (E) Summary of the physiological significance of Shu’s dispersion from Yb bodies. Following pre-Piwi-piRISC assembly, the complex moves to the outer mitochondrial membrane for Zuc-dependent maturation. The exit of Shu from Yb bodies is required for pre-Piwi-piRISC departure from the granule. On mitochondria, Gasz and Daed receive pre-Piwi-piRISC^29,63^. See also Figure S4.

Additionally, when Shu was fused to both EGFP and firefly luciferase (Luc), creating a construct with a molecular weight approximately equal to the combined molecular weights of Shu-EGFP and Shu-GBP-FLAG, the protein remained dispersed in the cytosol, confirming that the accumulation of Shu-EGFP and Shu-GBP-FLAG was not caused by an increase in molecular weight due to complex formation (Figures S4C and S4D).

We next examined the impact on the subcellular localization of Myc-Piwi. Myc-Piwi was cytosolic in the absence of Shu (Figures 1D, 4C, and S4E). When either Shu-EGFP or Shu-GBP-FLAG was expressed, Myc-Piwi localized to the nucleus. This suggests that Shu-EGFP alone or Shu-GBP-FLAG alone can rescue the effect of Shu loss on Myc-Piwi localization (Figures 4C and S4E). However, when Shu-EGFP and Shu-GBP-FLAG were coexpressed, the nuclear localization of Myc-Piwi was impaired, indicating a failure in piRISC biogenesis. Under this condition, Myc-Piwi tended to reside in Yb bodies along with Shu-EGFP and Shu-GBP-FLAG (Figures 4C and S4E). Thus, the forced accumulation of Shu in Yb bodies inhibits the departure of Piwi, presumably the Armi–pre-Piwi-piRISC complex, from Yb bodies.

To confirm this model, we next set up cellular environments similar to those described above, but with Zuc depleted, and examined the behavior of Armi under each condition. In Zuc-lacking OSCs, Armi accumulated on the mitochondrial surface, where the Armi–pre-Piwi-piRISC complex undergoes maturation^12,29^ (Figures 4D and S4F). In the additional absence of endogenous Shu, Armi tended to accumulate in Yb bodies (Figures 4D, S4F, and S4G), as predicted by the notion that pre-Piwi-piRISC assembly is necessary for the translocation of Armi from Yb bodies to the mitochondrial surface^12^. Ectopic expression of Shu-GBP-FLAG or Shu-EGFP under these conditions restored Armi’s steady-state localization (Figures 4D, S4F, and S4G), indicating that these ectopic Shu can rescue the loss of endogenous Shu if expressed separately. However, when they were coexpressed, Armi again accumulated in Yb bodies (Figures 4D, S4F, and S4G). The binding of Armi to RNA, which was previously reported as a trigger for the departure of the Armi–pre-Piwi-piRISC complex from Yb bodies^12^, remained largely unchanged by the forced accumulation of Shu in Yb bodies (Figure S4H). These findings support the intriguing idea that the rapid release of Shu from Yb bodies serves as an additional trigger for the detachment of the Armi–pre-Piwi-piRISC complex from Yb bodies (Figure 4E).

### The Armi point mutation corresponding to the nonobstructive azoospermia-associated mutation traps Shu in Yb bodies and disturbs Piwi-piRISC biogenesis

A recent human genetics study identified a mutation associated with nonobstructive azoospermia in the gene encoding MOV10L1, the human homolog of *Drosophila* Armi^46^. The importance of MOV10L1 in piRISC biogenesis in mice has already been confirmed^47,48^. The point mutation, 2447G to T, in the gene encoding MOV10L1 changed Ser816 in MOV10L1 to isoleucine^46^ (Figure 5A). However, detailed molecular analysis has not yet been conducted.

**Figure 5.**
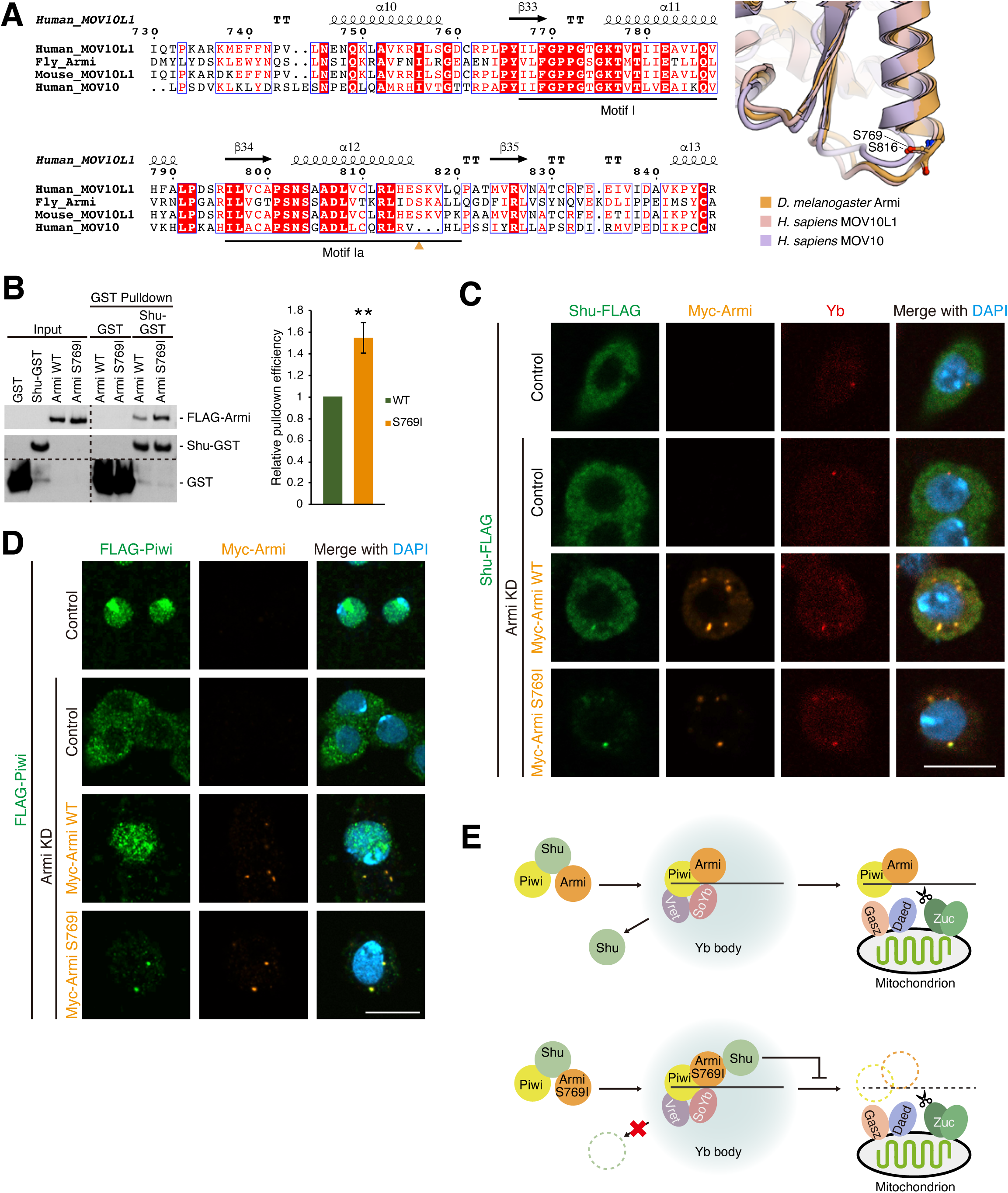
The Armi S769I mutant corresponding to azoospermia-associated mutant in human MOV10L1 represses dissociation of Shu from Yb bodies. (A) Left: sequence alignment of MOV10L1 homologs showing the position of the azoospermia-associated S816I mutation (orange triangle) in human MOV10L1. Helicase motifs^64^ are underlined. Right: superposition of predicted structures of MOV10L1 homologs. (B) Pull-down assay using recombinant proteins. The interaction between Shu and Armi was stabilized upon introduction of the S769I mutation in Armi. GST served as a negative control. The graph on the right shows quantified data. Error bars represent means ± SEM values from four independent experiments. **p < 0.01. (C) Rescue experiments revealed that the S769I mutation resulted in Shu accumulation in Yb bodies. Nuclei are shown in blue. Scale bar: 10 μm. (D) Rescue experiments showed that the S769I mutant resulted in accumulation of FLAG-tagged Piwi (FLAG-Piwi) in Yb bodies. Nuclei are shown in blue. Scale bar: 10 μm. (E) Summary of the effect of the S769I mutation in OSCs. Upper: wild-type Armi translocates from Yb bodies to mitochondria after Armi–pre-Piwi-piRISC assembly. Lower: the S769I mutation increases Armi’s affinity to Shu, trapping Shu in Yb bodies and inhibiting the Armi–pre-Piwi-piRISC departure from the granule. See also Figure S5.

We first sought to determine how the *Drosophila* Armi mutant, S769I, corresponding to the human MOV10L1 S816I mutant, influences the binding efficiency with Shu. To this end, we produced recombinant Armi WT and the S769I mutant and performed *in vitro* pull-down assays with GST-tagged Shu (Shu-GST) (Figure S5A). The Armi mutant bound to Shu-GST approximately 1.5 times more efficiently than the WT control (Figure 5B), suggesting that the point mutation has the effect of enhancing the Armi–Shu binding.

We next examined *in vivo* effects. Armi WT and the S769I mutant were individually expressed in OSCs lacking endogenous Armi. Simultaneously, Shu-FLAG was expressed in the cells, and its cellular localization was determined by immunofluorescence. In the absence of both endogenous and exogenous Armi, Shu-FLAG was scattered in the cytosol (Figures 5C and S5B). In the presence of Myc-tagged Armi (Myc-Armi) WT, Shu-FLAG began to appear in Yb bodies, although cytosolic signals were still dominant (Figures 5C and S5B). However, when Myc-Armi S769I mutant was alternatively expressed, Shu-FLAG strongly accumulated in Yb bodies. The RNA unwinding activity of Armi was minimally affected by the mutation (Figure S5C).

We then examined how this situation impacts the subcellular localization of Piwi. FLAG-tagged Piwi (FLAG-Piwi) was detected in the nucleus in OSCs expressing Myc-Armi WT, as expected, indicating that Piwi-piRISC production was successful (Figures 5D and S5D). However, when the Myc-Armi S769I mutant was alternatively expressed, the Piwi signals appeared mostly in Yb bodies, while the nuclear signals were nearly negligible. These results indicate that the Armi S769I mutant disturbs the production of Piwi-piRISC, likely by hindering the proper release of the Armi–pre-Piwi-piRISC complex from Yb bodies (Figure 5E). This impaired release mirrors the defects observed under conditions of forced Shu self-association, further supporting the notion that rapid ejection of Shu from Yb bodies is critical for Piwi-piRISC biogenesis and the functionality of Piwi in OSCs.

## Discussion

This study reveals that Shu localizes to Yb bodies in an Armi-dependent manner to deposit Piwi in the bodies but quickly returns to the cytosol through the electrostatic repulsion of its N-terminal acidic residues. This exit of Shu from the granules is necessary for Armi to transfer pre-Piwi-piRISC from Yb bodies to the mitochondrial surface, the place of piRISC maturation. A point mutation in Armi, analogous to a mutation in human MOV10L1 associated with nonobstructive azoospermia, disrupts this process by stabilizing the Armi–Shu interaction in Yb bodies, thereby inhibiting Piwi-piRISC production.

Using Yb bodies and Shu as models, we demonstrated that the repulsive forces of client proteins contribute to the transient nature of their granule localization. Our findings expand the understanding of electrostatic forces in regulating the granule localization of proteins. Previous studies showed that artificially increased repulsion of a scaffold protein, the DNA-binding protein of 43 kDa (also known as TDP-43), inhibits pathological amyloid formation^49^, and that the salt concentration dependence of LLPS in scaffold proteins like bovine serum albumin (BSA) and DDX4 is also mediated by electrostatic repulsion^50,51^. Although previous reports have indicated that electrostatic repulsion among scaffold proteins and/or client proteins is related to the regulation of scaffold LLPS, none have analyzed it at the molecular level in as much detail as this study. The physiological importance of the release of client protein Shu from Yb bodies is experimentally validated by our study. We anticipate that the importance of protein dispersion will be discovered in other cellular processes.

This study and the previous study^12^ identified two requirements for the departure of Piwi-pre-piRISC from Yb bodies to mitochondria: Shu release and Armi’s RNA-binding activity. This multilayered regulation likely ensures that only properly assembled Piwi-pre-piRISC complexes are allowed to exit Yb bodies. In particular, the dependence on Shu release could serve as a checkpoint, ensuring that Piwi is transferred to SoYb/Vret heterodimers before leaving the granules. Further investigation into the molecular role of SoYb/Vret during Piwi-pre-piRISC assembly is warranted.

We found that the mutation associated with human infertility causes aberrant localization of Shu in OSCs, underscoring the interspecies conservation of the importance of electrostatic control of Shu localization. FKBP6, the mouse counterpart of Shu, is reported to disperse in the cytosol and not accumulate in pi-bodies, which are mouse germ granules involved in the embryonic piRNA pathway^52^. Loss of FKBP6 causes the mouse nuclear Piwi protein MIWI2 to fail to load piRNAs and to disperse in the cytosol^52^. Lack of expression of human FKBP6 is also associated with oligozoospermia^53^. These observations suggest that the role of Shu in the piRNA pathway, as well as the transient localization in droplets, is conserved across species. Notably, these previous studies proposed that Shu/FKBP6 functions downstream of known piRNA factors, particularly in the loading of mature piRNAs on Piwi proteins, in individual *Drosophila* and mice^31,52^. Our present study supports the idea that Shu is one of the most upstream factors in Piwi-piRISC biogenesis occurring in OSCs.

Defects in the piRNA pathway result in infertility across model animals. Mutations in human orthologs of piRNA pathway genes are also found in male infertile patients through genomic surveillance. However, the functional impact of single amino acid substitutions remains poorly understood. Recently, oligozoospermia-associated missense mutations in HIWI and PNLDC1, proteins whose mouse orthologs are critical in the postnatal piRNA pathway^54,55^, were analyzed biochemically^56,57^. Our study provides critical biochemical insight into the effect of the MOV10L1 S816I mutation, which is linked to azoospermia through its potential disruption of embryonic piRNA biogenesis, offering a potential target for therapeutic intervention.

The interaction between Hsp83 and Shu was not detected in OSCs (Figure S1C). This result is consistent with previous research showing that the introduction of the K304A mutation in Shu, which is expected to weaken its interaction with Hsp83, barely affects the subcellular localization of Piwi in *Drosophila* ovaries^31^. Thus, the Shu– Hsp83 interaction may not be required for piRNA loading to Piwi. In contrast, the subcellular localization of Aubergine and Ago3, germline-specific PIWI proteins in *Drosophila*, is altered by the K304 mutation^31^. The loading of RNAs onto Argonaute proteins is enhanced by Hsp83/Hsp90 and co-chaperones in various model organisms, including *Arabidopsis*^58–62^, implying that co-chaperones have assisted in the loading of small RNAs to Argonaute long before the acquisition of piRNAs. In theory, Armi/MOV10L1 could bind directly to PIWI and transport it to cytoplasmic granules. However, Armi/MOV10L1 has a second function in piRISC biogenesis, namely, to transport pre-PIWI-piRISC to mitochondria for Zuc-dependent maturation. Therefore, it is likely that Shu/FKBP6 was employed as a marker for PIWI proteins that are ready for loading piRNA intermediates and is recognized by Armi/MOV10L1 as tags on PIWI directing transport to the cytosolic granules. Following the acquisition of this mechanism, *Drosophila* Piwi may have lost its dependence on the Shu–Hsp83 interaction for RNA loading.

## Resource Availability

### Lead contact

Requests for further information and resources should be directed to and will be fulfilled by the lead contact, Mikiko C. Siomi.

## Materials availability

Plasmids and antibodies produced in this study are available upon request to the lead author.

## Data and code availability

This paper does not report original code. All original data and additional information required to reanalyze the results presented in this paper are available from the lead contact upon reasonable request.

## Acknowledgements

We are grateful to J. Brennecke, K. Sato, and K. Nakayama for sharing vectors. We are also thankful to Y. Tomoe for technical support. We acknowledge Y. Hara and R. Hayashi (Carl Zeiss Co., Ltd.) for technical assistance in microscopy. We also thank other members of the Siomi laboratory for discussions and comments on the manuscript. This work was supported by JSPS KAKENHI grant numbers 24K09376 (to S.H.) and 19H05466 (to M.C.S.).

## Author Contributions

S.H. performed experiments with support from A.F.. S.H. and M.C.S. designed the experiments and wrote the manuscript. M.C.S. supervised all the research.

## Declaration of Generative AI and AI-assisted Technologies in the Writing Process

During the preparation of this work the authors used ChatGPT and Perplexity for proofreading. After using these tools, the authors reviewed and edited the content as needed and take full responsibility for the content of the published article.

## Supplementary Figure Legends

**Figure S1. Related to Figure 1.**
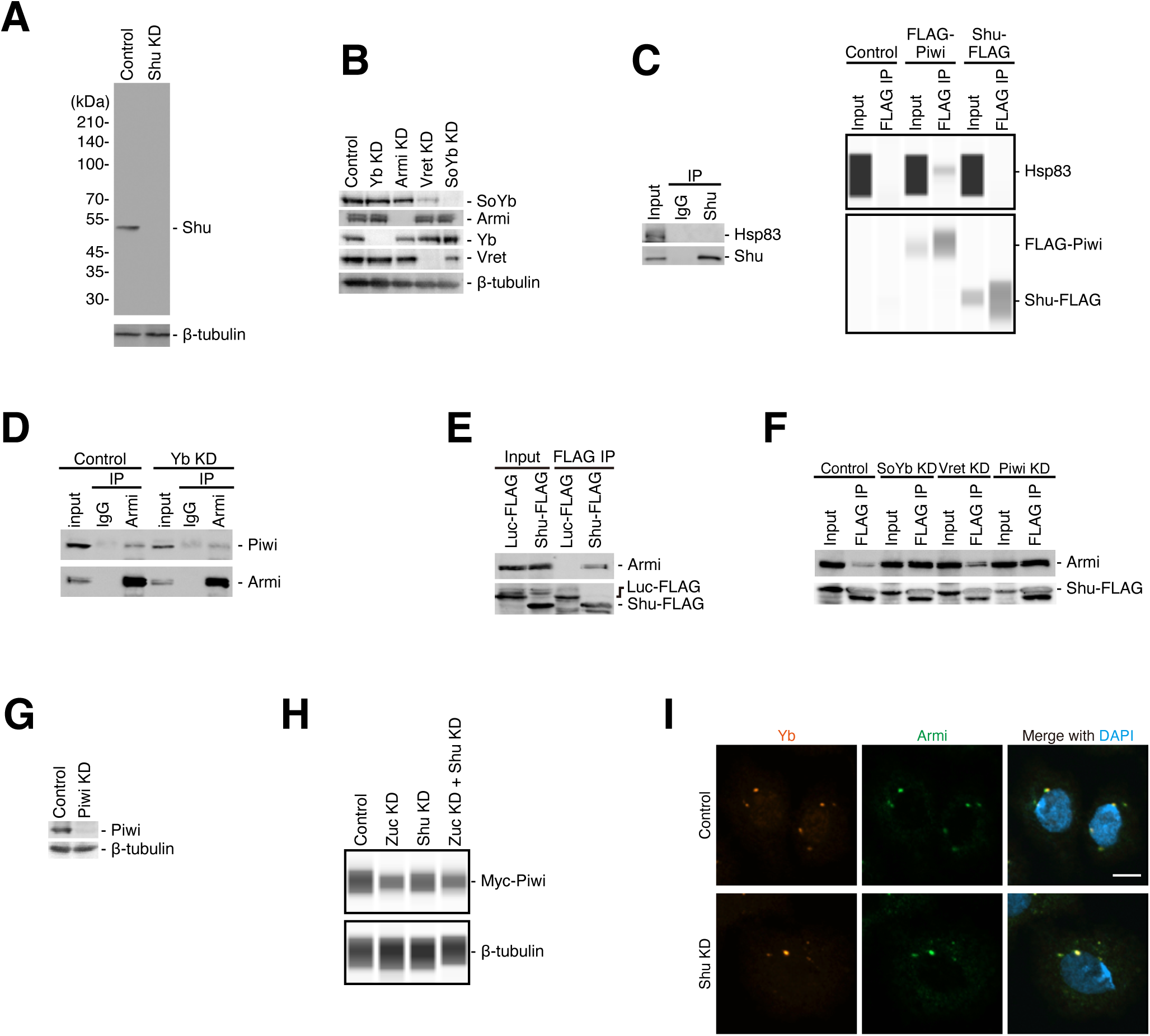
(A) Western blotting demonstrating the specificity of the anti-Shu polyclonal antibody produced in this study. β-tubulin is shown as a loading control. (B) The efficiency of knockdown (KD) of Yb body components. SoYb and Vret stabilize each other as reported previously^8^. β-tubulin serves as a loading control. (C) Hsp83 was not detected in the Shu (left) or Shu-FLAG (right) complex immunoisolated from OSCs. FLAG-Piwi was used as a positive control^65^. Non-immune IgG was used as a negative control. (D) The interaction between Armi and Piwi did not require Yb (i.e., Yb bodies). Non-immune IgG was used as a negative control. (E) Shu-FLAG complex immunoisolated from OSCs using anti-FLAG antibodies contains Armi. Luc-FLAG was used as a negative control. (F) Shu-FLAG and Armi remained bound to each other upon SoYb, Vret, or Piwi depletion. (G) The KD efficiency of Piwi. β-tubulin is shown as a loading control. (H) Western blotting confirming Myc-Piwi expression in the cells shown in Figure 1D. β-tubulin is a loading control. (I) Armi still localized in Yb bodies upon Shu depletion. Nuclei are shown in blue. Scale bar: 5 μm.

**Figure S2. Related to Figure 2.**
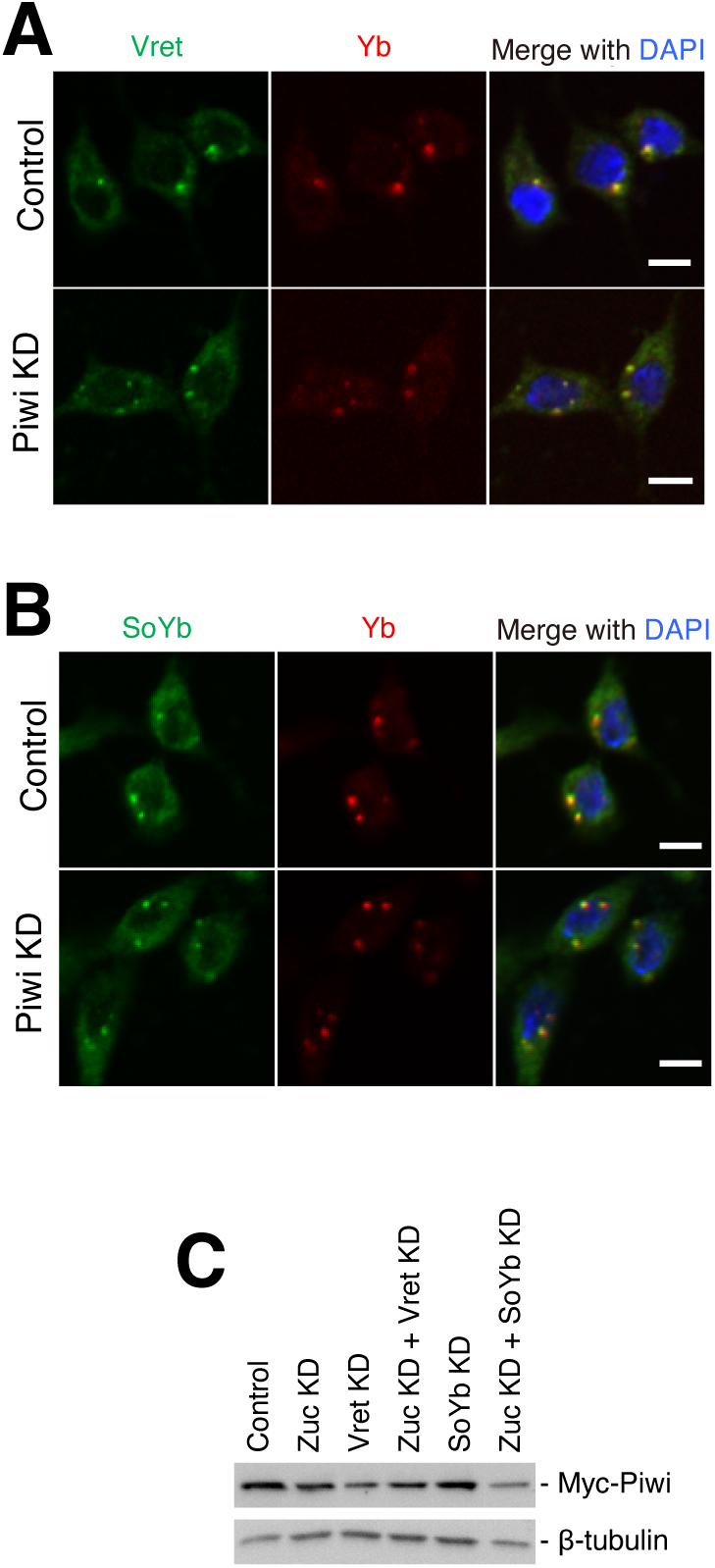
(A, B) Vret (A) and SoYb (B) remained localized in Yb bodies upon depletion of Piwi. Nuclei are shown in blue. Scale bar: 5 μm. (C) Western blotting showing Myc-Piwi is expressed in the cells shown in Figure 2E. β-tubulin is a loading control.

**Figure S3. Related to Figure 3.**
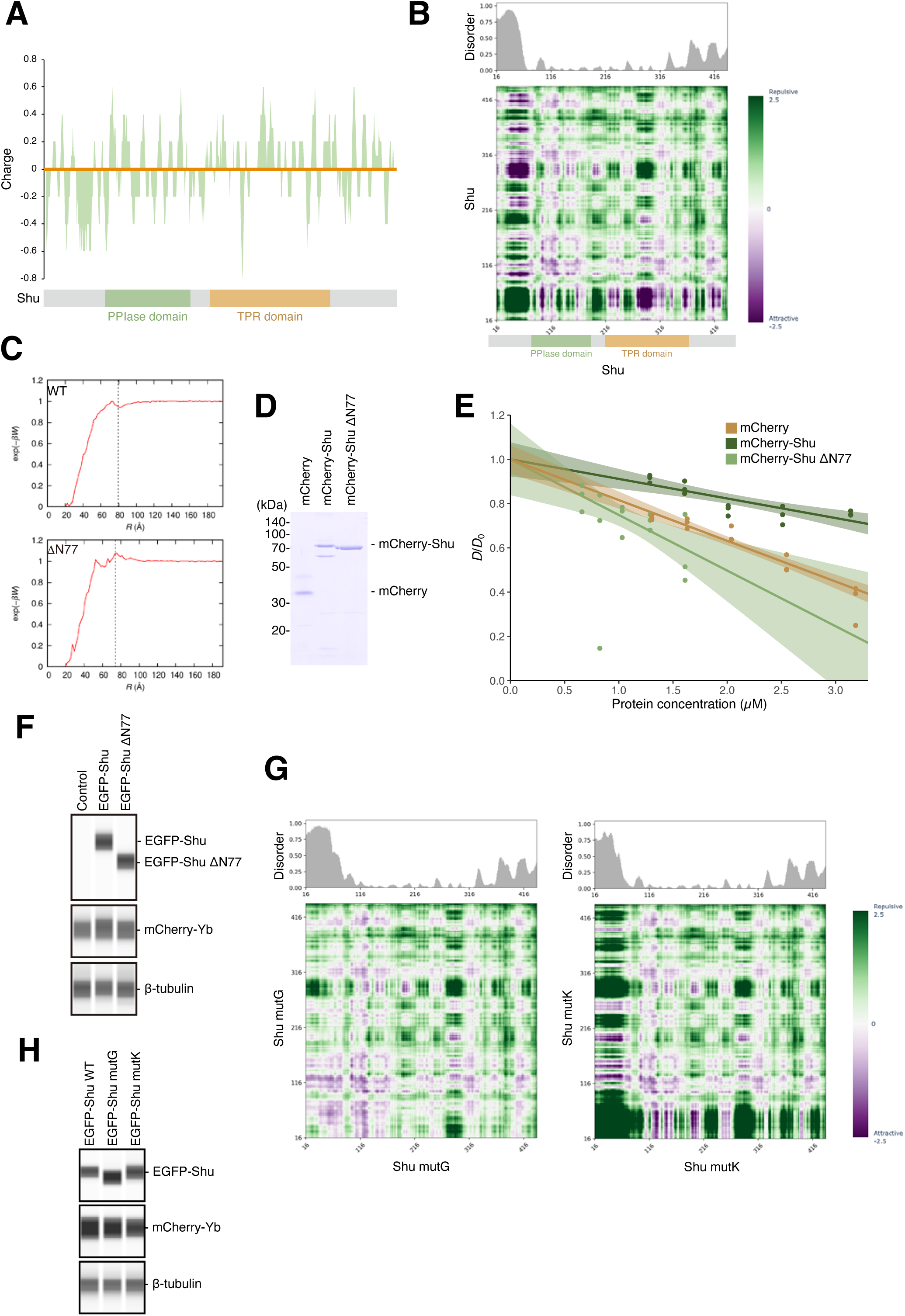
(A) Charge distribution of Shu. (B) Upper: the N-terminus of Shu is predicted to be disordered. Lower: predicted interaction map of Shu self-interaction. Strong repulsion between N-terminal disordered residues is shown. The interaction map of the structured region is also shown for reference, but protein folding was not considered during prediction. (C) The potential of mean force (PMF), denoted as *W*, was predicted, assuming rigid structures. The corresponding Boltzmann factors, represented as exp(-*βW*) (where *β* =1/(*k*_B_*T*), with *k*_B_ as the Boltzmann constant and *T* as the temperature) were plotted as a function of interprotein distances. For Shu lacking N-terminal 77 residues (ΔN77), the highest probability density was predicted at a 7.4 nm distance, similar to the diameter of the equivalent sphere of protein (dashed line). PMF of wild-type Shu is also predicted without considering N-terminal flexibility and is shown for reference. (D) CBB-stained recombinant mCherry-tagged proteins after SDS-PAGE. (E) Diffusion coefficients (*D*) were plotted against protein concentrations and normalized to intercepts (*D*_0_). Confidence intervals of regression lines were shown as shades. Slopes of the regression lines represent *k_D_* values in Figure 3B. (F) Western blotting confirming EGFP-Shu and mCherry-Yb expression in the cells shown in Figure 3E. β-tubulin is shown as a loading control. (G) Predicted interaction map of self-interaction of the Shu mutants. The maps of structured regions are also shown for reference. Mutated regions were still predicted to be disordered as shown on the top. (H) Western blotting showing EGFP-Shu and mCherry-Yb were expressed in the cells shown in Figure 3G. β-tubulin is a loading control.

**Figure S4. Related to Figure 4.**
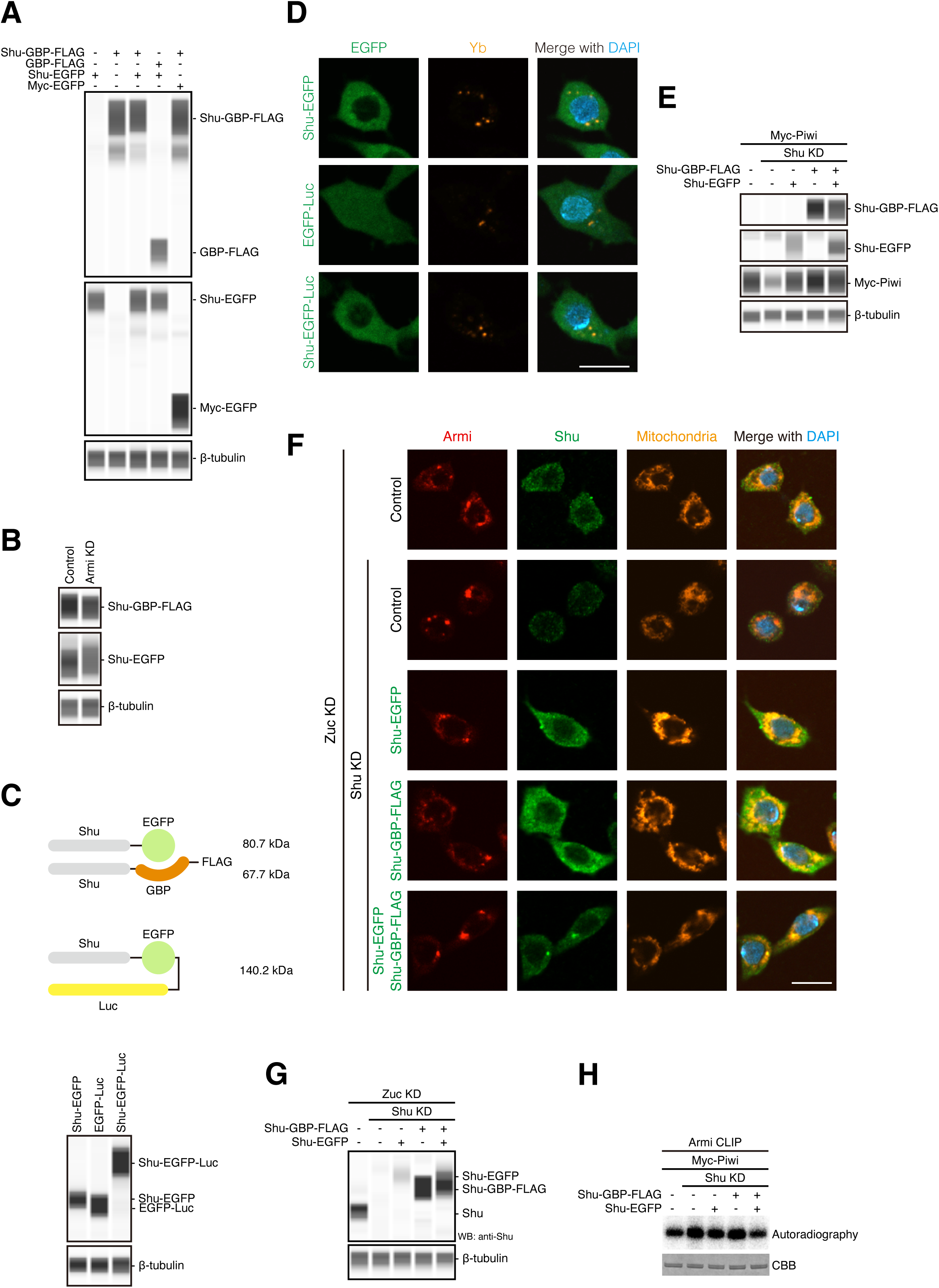
(A, B) Western blotting showing tagged Shu was expressed in the cells shown in Figures 4A (A) and 4B (B). β-tubulin is shown as a loading control. (C) Upper: Luc was fused to Shu-EGFP to bring its molecular weight approximately equal to the combined molecular weights of Shu-EGFP and Shu-GBP-FLAG. Lower: western blotting showing expression of EGFP and Luc-tagged Shu (Shu-EGFP-Luc) in OSCs. β-tubulin serves as a loading control. (D) Shu-EGFP-Luc barely accumulated in Yb bodies. Nuclei are shown in blue. Scale bar: 10 μm. (E) Western blotting confirming tagged Shu expression in the cells shown in Figures 4C. β-tubulin is shown as a loading control. (F) Rescue experiments showing that forced Shu accumulation in Yb bodies inhibited Armi mitochondrial localization in Zuc-depleted cells. Part of the mitochondria clustered near Yb bodies, as previously reported^21,66^. Nuclei are shown in blue. Scale bar: 10 μm. (G) Western blotting showing tagged Shu proteins were expressed in the cells shown in Figures 4D and S4F. β-tubulin is a loading control. (H) Crosslinking-immunoprecipitation showing that Armi interacts with RNA in the cells shown in Figure 4C.

**Figure S5. Related to Figure 5.**
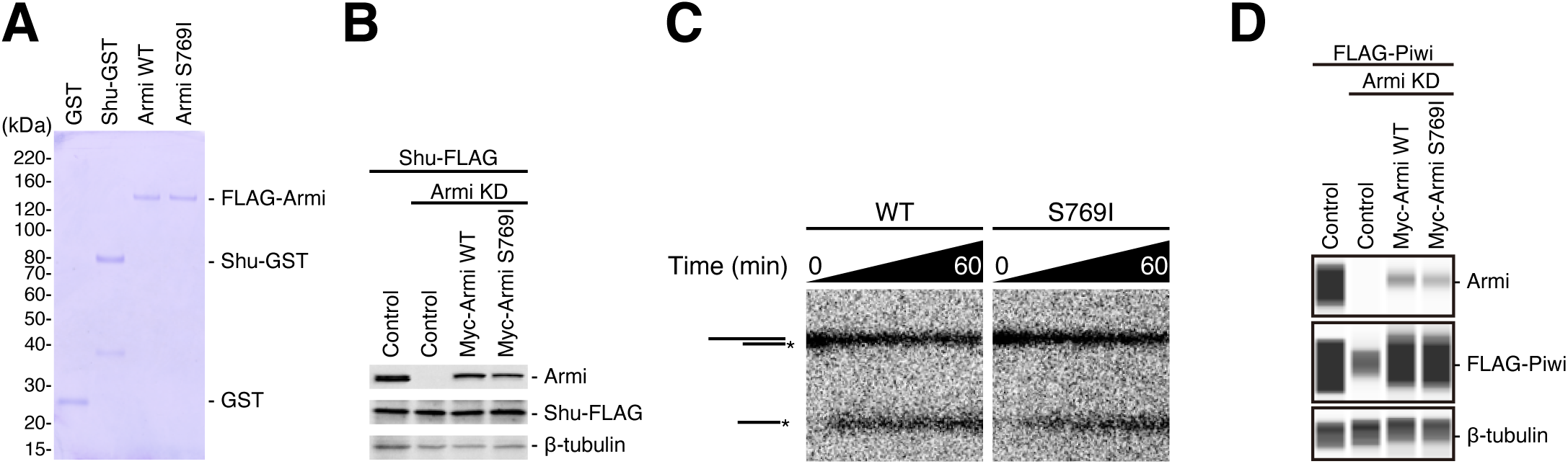
(A) CBB-stained recombinant proteins used in the assays shown in Figures 5B and S5C. (B) Western blotting showing tagged proteins were expressed in the cells shown in Figure 5C. β-tubulin is shown as a loading control. (C) Unwinding activity of recombinant Armi wild-type (left) and S769I mutant (right). RNA duplex with 5′ overhang was incubated with FLAG-tagged Armi (FLAG-Armi) for varying time intervals, then separated by non-denaturing PAGE. The asterisks indicate the ^32^P label positions. (D) Western blotting showing tagged proteins were expressed in the cells shown in Figure 5D. β-tubulin is shown as a loading control.

## STAR★Methods

### Key resource table

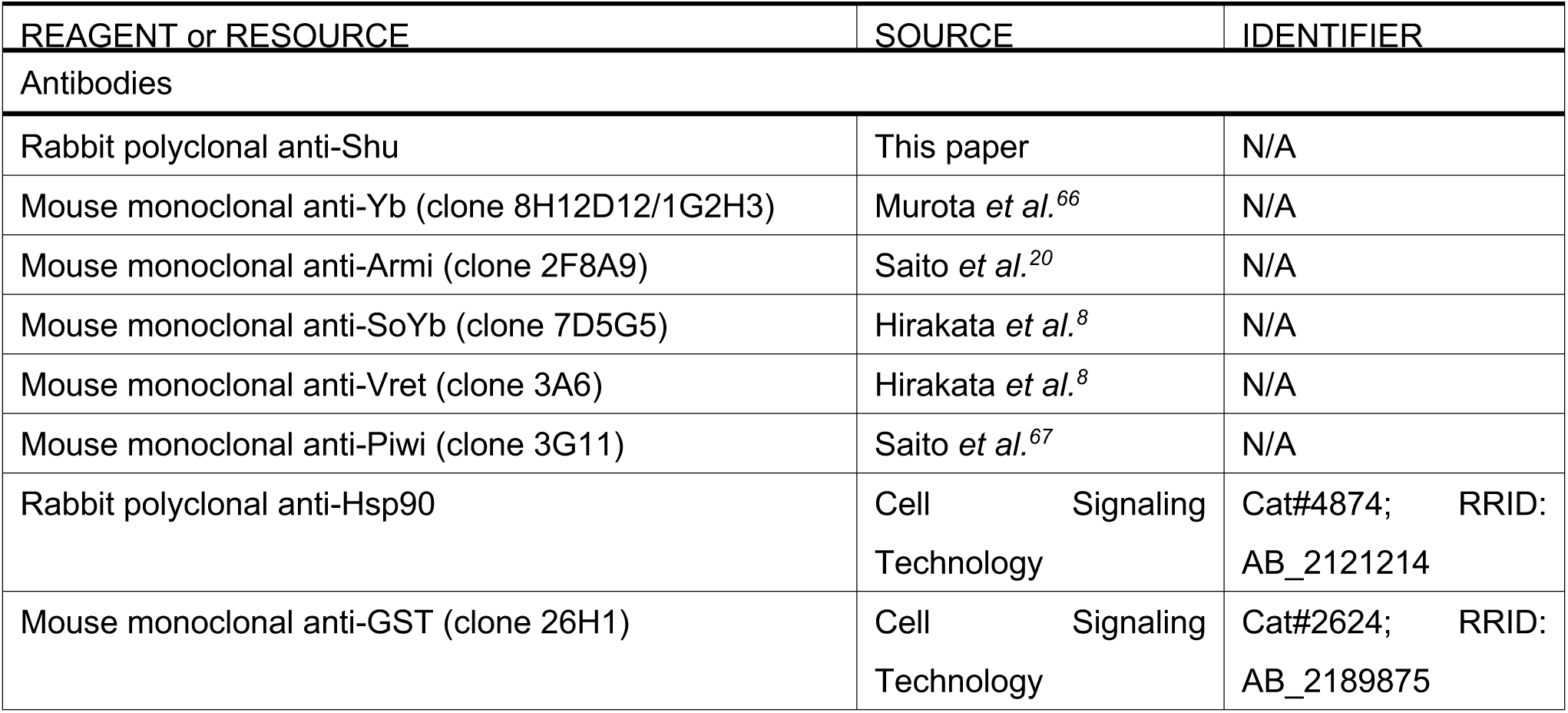
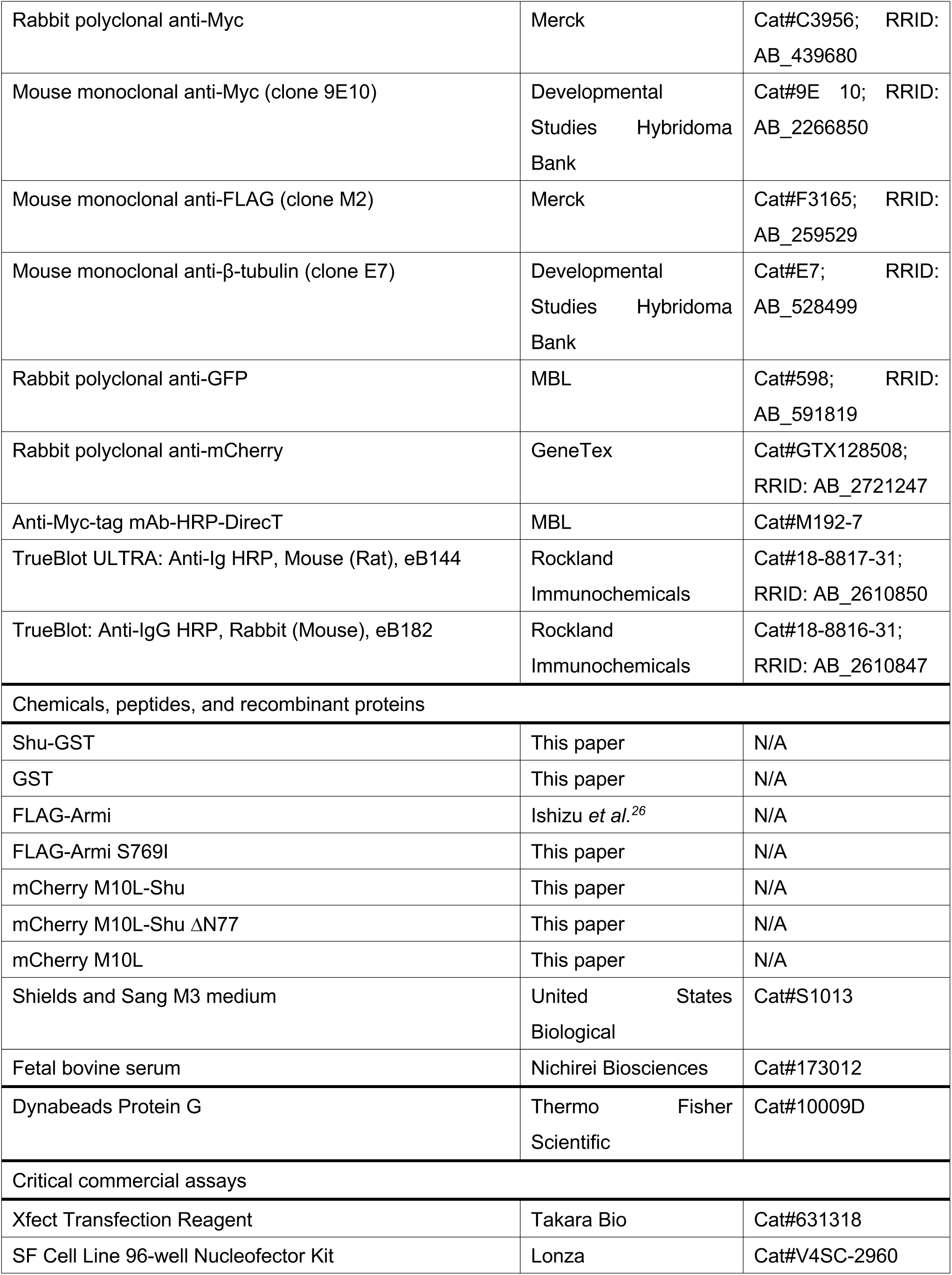
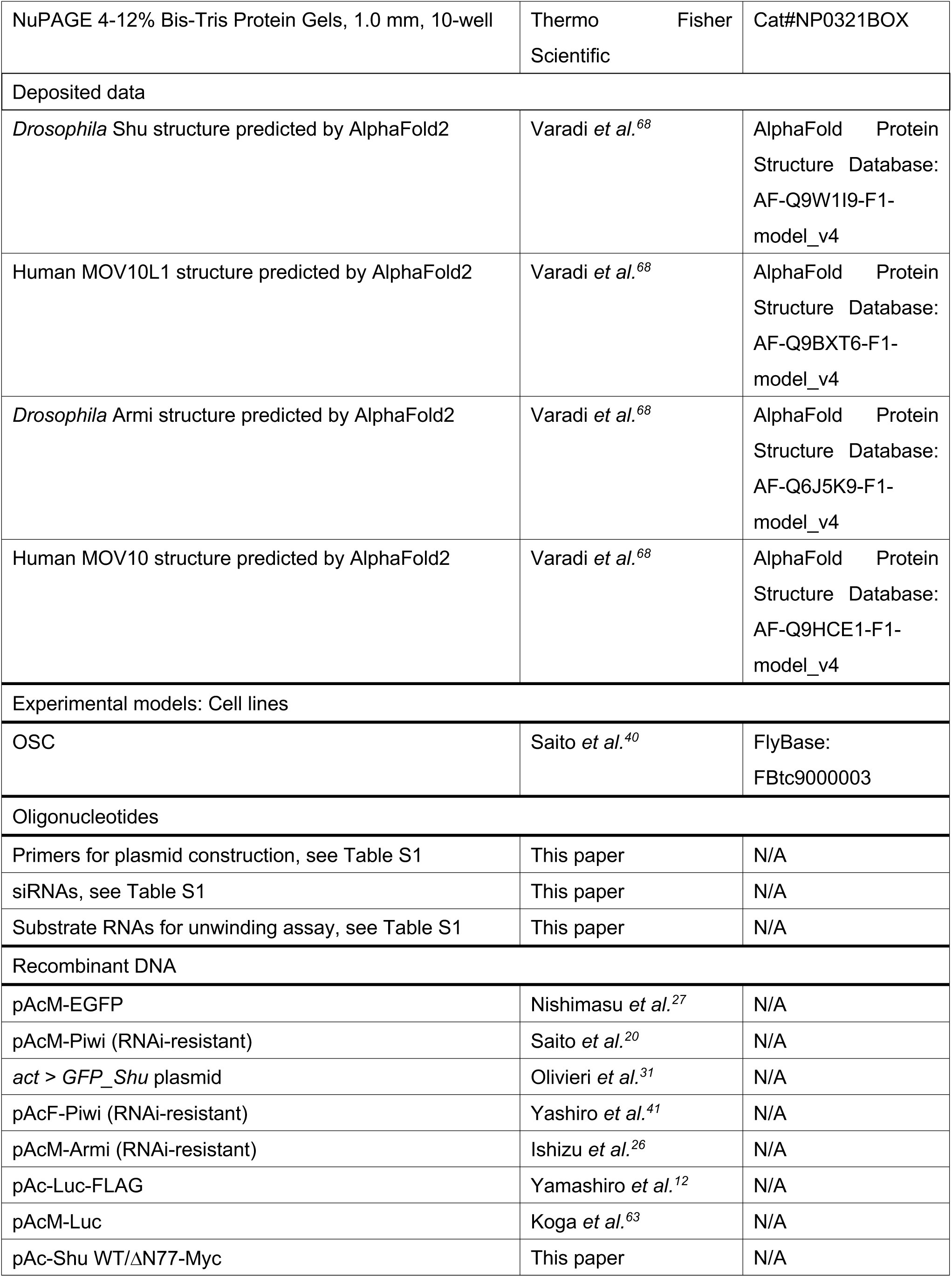
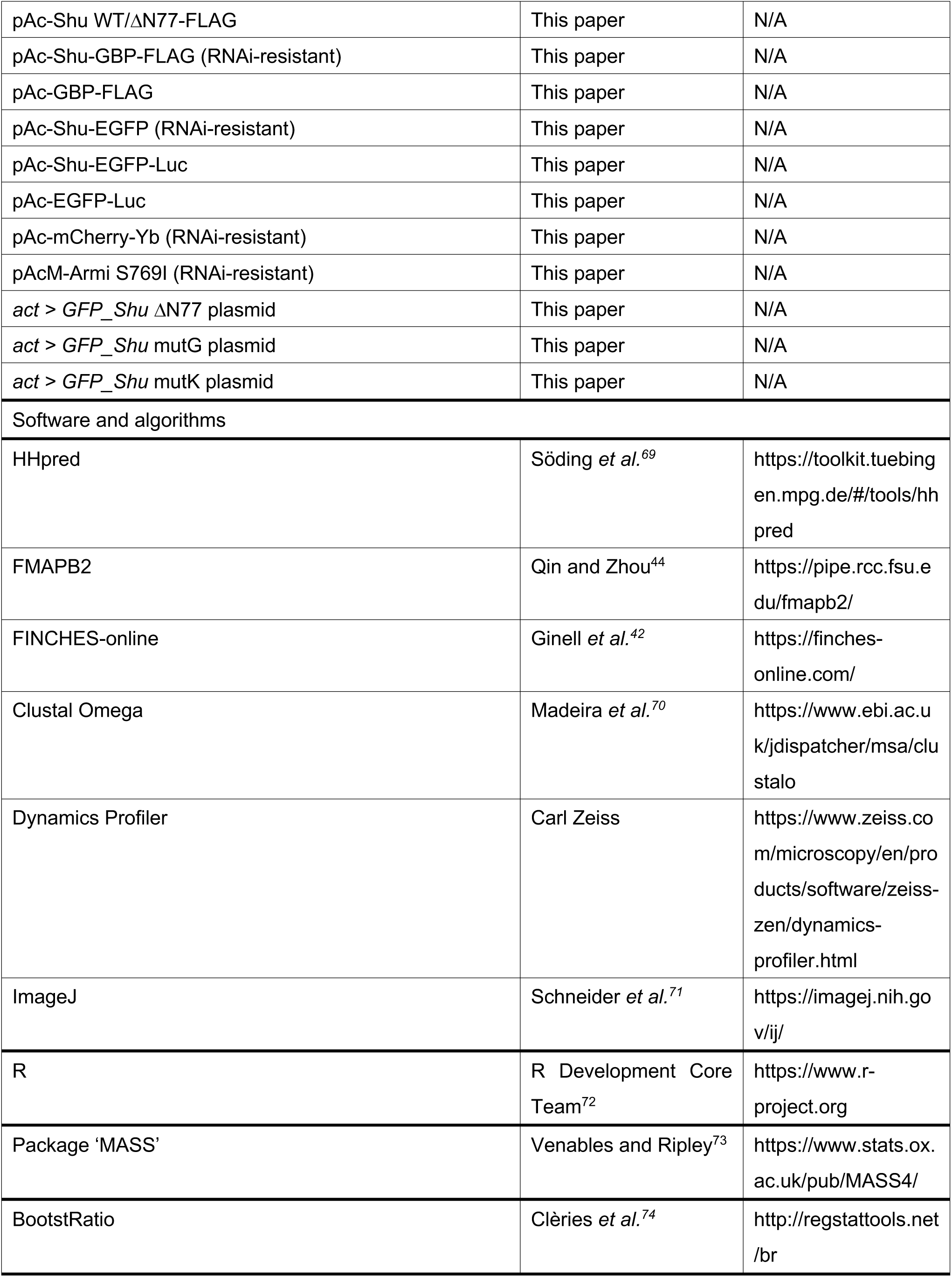
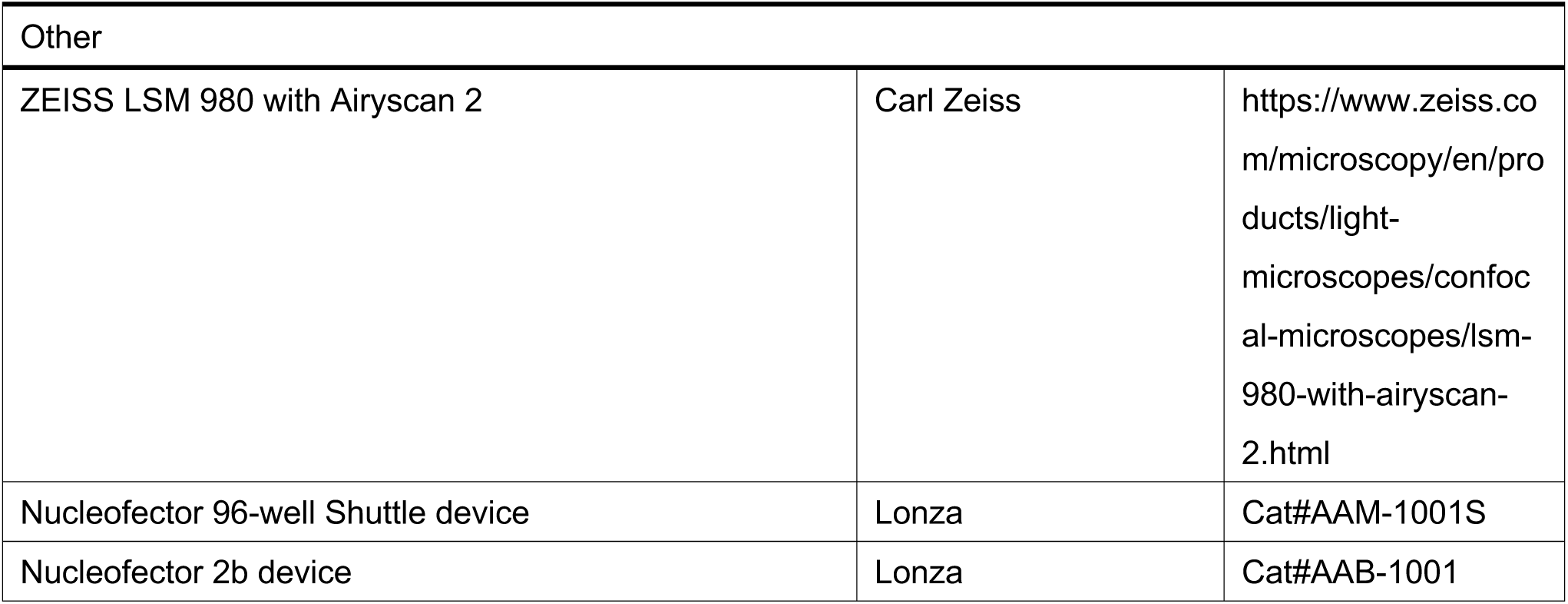

### Experimental model and study participant details

#### Cell lines

OSC^40^ was cultured at 26°C in Shields and Sang M3 medium (United States Biological) supplemented with 10% fetal bovine serum (FBS; Nichirei Biosciences), 10 mU/mL insulin, 0.6 mg/mL glutathione, 10% fly extract, and 1x Antibiotic-Antimycotic (Thermo Fisher Scientific). Fly extract was prepared by heat-inactivating a fly homogenate (0.2 g/mL adult Oregon R flies in Shields and Sang M3 medium supplemented with 10% FBS) at 60°C for 5 minutes, followed by removal of precipitates.

### Method details

#### Plasmid construction

The expression vectors of Myc-Piwi^20^, Myc-EGFP^27^, EGFP-Shu^31^, FLAG-Piwi^41^, Myc-Armi^26^, FLAG-Armi^26^, Luc-FLAG^12^, and Myc-Luc^63^ were described previously.

To generate the Shu-Myc expression vector, a DNA fragment encoding Shu was amplified by PCR from OSC cDNA and cloned into a PCR-linearized pAcM vector^40^ using NEBuilder HiFi DNA Assembly Master Mix (New England Biolabs). For the Shu-FLAG expression vector, the Myc-encoding region was removed from the Shu-Myc vector by inverse PCR, and a 3xFLAG sequence, amplified from the FLAG-Armi vector, was inserted. To construct the Shu-GBP-FLAG vector, a DNA fragment encoding GBP (GFP-nanobody) was amplified from the pGEX6P1-GFP-Nanobody [a gift from Kazuhisa Nakayama (Addgene plasmid # 61838; http://n2t.net/addgene:61838; RRID:Addgene_61838)]^75^ and cloned into a PCR-linearized Shu-FLAG vector. Synonymous substitutions were introduced into the regions encoding Shu in both Shu-EGFP (a gift from Julius Brennecke) and Shu-GBP-FLAG vectors by inverse PCR to render them RNAi-resistant. The GBP-FLAG expression vector was created by deleting the Shu region from the Shu-GBP-FLAG vector via inverse PCR. To construct expression vectors of Shu-EGFP-Luc and EGFP-Luc, the Myc-encoding region was removed from the Myc-Luc vector by inverse PCR, and DNA fragments encoding Shu and EGFP were amplified from the Shu-Myc and Myc-EGFP vectors, respectively, and inserted. For the mCherry-Yb expression vector, the EGFP-encoding region was removed from the EGFP-Yb vector (Hirakata et al., 2019) by inverse PCR, and an mCherry-encoding fragment, amplified from the Myc-mCherry vector (a gift from Kaoru Sato), was inserted. For the *E. coli* expression vector of Shu-GST, the region encoding the N-terminal GST tag was deleted from pGEX6p-1 (Cytiva) by inverse PCR, and DNA fragments encoding Shu and GST were amplified from the Shu-FLAG vector and pGEX6p-1, respectively, and inserted. To construct the *E. coli* expression vectors of mCherry-Shu-GST and mCherry-GST, a DNA fragment of the mCherry CDS, amplified from the mCherry-Yb vector, was inserted into a PCR-linearized Shu-GST vector. By inverse PCR, the alternative start codon of mCherry was substituted^76^, and the N-terminal sequences of mCherry-Shu-GST and mCherry-GST were equalized.

Expression vectors of Myc-Armi S769I, FLAG-Armi S769I, Shu ΔN77-Myc, Shu ΔN77-FLAG, EGFP-Shu ΔN77, and mCherry-Shu ΔN77-GST were generated by inverse PCR using the wild-type vectors as templates. For the EGFP-Shu mutK expression vector, site-directed mutagenesis was performed with NEBuilder HiFi DNA Assembly Master Mix using the EGFP-Shu expression vector as the template. For the EGFP-Shu mutG expression vector, eleven acidic residues were substituted with Gly by inverse PCR, followed by mutation of the remaining nine acidic residues via site-directed mutagenesis using NEBuilder HiFi DNA Assembly Master Mix.

The primer sequences used for these constructions are listed in Table S1.

#### RNAi and transfection with plasmids

RNAi was performed using a Nucleofector 96-well Shuttle device (program DG-150) with SF Cell Line 96-well Nucleofector Kit (Lonza Bioscience). Plasmid transfections were carried out using Xfect Transfection Reagent (TaKaRa Bio) following the manufacturer’s protocol. Co-transfection of plasmids and siRNAs was performed using a Nucleofector 2b device (Lonza Bioscience) or with a combination of Xfect and the Nucleofector 96-well Shuttle device. For transfection using the Nucleofector 2b device, cells were suspended in transfection buffer [180 mM sodium phosphate buffer for Church and Gilbert hybridization (pH 7.2)^77^, 5 mM KCl, 15 mM MgCl_2_, 50 mM D-mannitol]^8^ with plasmids and siRNAs, and electroporation was performed in a 2-mm gap electroporation cuvette using program N-020.

In RNAi experiments, cells were re-transfected with siRNAs 48 hours after the initial transfection. For exogenous gene expression in knockdown cells, plasmids and siRNAs were co-transfected 48 hours after the first siRNA transfection. Cells were harvested 24 or 48 hours after the final transfection. The siRNA sequences used in this study are listed in Table S1.

#### Immunofluorescence

OSCs were fixed with 4% formaldehyde in phosphate-buffered saline (PBS) and permeabilized with 0.1% Triton X-100 in PBS. Immunostaining was carried out using antibodies diluted in 3% BSA in PBS, with PBS washing following each treatment. Cells were mounted with VECTASHIELD Mounting Medium with DAPI (Vector Laboratories) and observed using an LSM 980 laser scanning confocal microscope (Carl Zeiss). Fluorescence from EGFP-or mCherry-tagged proteins was also detected in relevant experiments. For mitochondrial staining, cells were incubated with MitoTracker Orange CMXRos (Thermo Fisher Scientific) at 1:2,000 dilution in OSC medium for 20 minutes prior to cell fixation. Unless otherwise denoted, more than 15 cells were examined, and representative images are presented.

The following primary antibodies were used: anti-Yb mouse monoclonal antibody (1:250 dilution)^66^, anti-Armi mouse monoclonal antibody (1:200 dilution)^20^, anti-Vret mouse monoclonal antibody (1:300 dilution)^8^, anti-SoYb mouse monoclonal antibody (1:500 dilution)^8^, anti-Shu rabbit polyclonal antibody (1:4.8 dilution; this study), anti-Myc rabbit polyclonal antibody (1:500 dilution; Merck), and anti-FLAG mouse monoclonal antibody (1:1,000 dilution; Merck). The polyclonal Shu antibody was produced by Eurofin using a synthetic peptide corresponding to Shu amino acids 31-44 as the antigen. Alexa Fluor 488/555/633-conjugated anti-rabbit IgG/mouse IgG1/mouse IgG2b/mouse IgG2a antibodies (Thermo Fisher Scientific) were used at 1:1000 dilution as secondary antibodies.

#### Manual western blotting

Proteins were separated on sodium dodecyl sulfate (SDS) polyacrylamide gels using either Laemmli electrophoresis running buffer or AllView PAGE Buffer (BioDynamics Laboratory), followed by transfer to nitrocellulose membranes. The membranes were incubated sequentially with 5% skimmed milk in 0.1% Tween-20 in PBS (T-PBS), primary antibodies diluted in T-PBS, and secondary antibodies diluted in T-PBS, with extensive washing in T-PBS after each step. Detection was carried out using Clarity Western ECL Substrate (Bio-Rad), and chemiluminescence images were captured using a ChemiDoc XRS Plus System (Bio-Rad).

The primary antibodies used were as follows: anti-Piwi mouse monoclonal antibody (1:1,000 dilution)^67^, anti-Armi mouse monoclonal antibody (1:1,000 dilution)^20^, anti-Vret mouse monoclonal antibody (1:1,000 dilution)^8^, anti-SoYb mouse monoclonal antibody (1:1,000 dilution)^8^, anti-Yb mouse monoclonal antibody (1:1,000 dilution)^66^, anti-Shu rabbit polyclonal antibody (1:100 dilution; this study), anti-Hsp90 rabbit polyclonal antibody (1:1,000 dilution; Cell Signaling Technology), anti-GST mouse monoclonal antibody (1:1,000 dilution; Cell Signaling Technology), anti-Myc mouse monoclonal antibody (1:1,000 dilution; Developmental Studies Hybridoma Bank), anti-FLAG mouse monoclonal antibody (1:10,000 dilution; Merck), anti-β-tubulin mouse monoclonal antibody (1:1,000 dilution; Developmental Studies Hybridoma Bank), and peroxidase-conjugated anti-Myc antibody (1:1,000 dilution; MBL). Secondary antibodies used in this study were peroxidase-conjugated anti-mouse IgG antibody (1:5,000 dilution; Cappel), peroxidase-conjugated anti-rabbit IgG antibody (1:1,000 dilution; Cell Signaling Technology), and peroxidase-conjugated TrueBlot antibodies against mouse and rabbit IgG (1:1,000 dilution; Rockland Immunochemicals).

#### Automated western blotting

Automated western blotting was performed using Abby (ProteinSimple) according to the manufacturer’s protocol. OSCs were lysed in Laemmli SDS sample buffer containing 100 mM dithiothreitol (DTT) and boiled at 95°C for 5 minutes. Prior to capillary electrophoresis using the 12-230 kDa Separation Module (ProteinSimple), 5x Fluorescent Master Mix (ProteinSimple) was added to the lysate. Blocking and dilution of all primary antibodies were performed using 5% skimmed milk in T-PBS, with precipitates removed. Detection was performed using the Anti-Mouse Detection Module, Anti-Rabbit Detection Module, and RePlex Module (ProteinSimple).

The primary antibodies used for automated western blotting were anti-β-tubulin mouse monoclonal antibody (1:25 dilution; Developmental Studies Hybridoma Bank), anti-Myc mouse monoclonal antibody (1:50 dilution; Developmental Studies Hybridoma Bank), anti-FLAG mouse monoclonal antibody (1:1,250 dilution; Merck), anti-GFP rabbit polyclonal antibody (1:50 dilution; MBL), anti-mCherry rabbit polyclonal antibody (1:50 dilution; GeneTex), anti-Hsp90 rabbit polyclonal antibody (1:50 dilution; Cell Signaling Technology), anti-Armi mouse monoclonal antibody (1:50 dilution)^20^, and anti-Shu rabbit polyclonal antibody (1:100 dilution; this study).

#### Immunoprecipitation

OSCs were lysed in binding buffer [50 mM Tris-HCl (pH 8.0), 150 mM KOAc, 5 mM Mg(OAc)_2_, 5 mM DTT, 0.1% NP-40, 2 µg/mL leupeptin, 2 µg/mL pepstatin A, 0.5% aprotinin, 10% glycerol] and centrifuged to remove insoluble debris. The lysates were then incubated with anti-Shu (this study), anti-Piwi^67^, anti-Armi^20^, or anti-FLAG (Merck) antibodies bound to Dynabeads Protein G (Thermo Fisher Scientific) at 4°C for 2 hours. Normal mouse IgG (FUJIFILM Wako Pure Chemical) was used as a negative control. The beads were washed five times with binding buffer, and proteins were eluted in 2x Laemmli SDS sample buffer without DTT. The eluates were then mixed with DTT to a final concentration of 100 mM, boiled, and used for manual or automated western blotting (see above).

For the analysis of the interaction between Shu and Armi, a buffer used in a previous study was modified and used^31^. OSCs were lysed in lysis buffer [10 mM Tris-HCl (pH 7.5), 150 mM NaCl, 0.5 mM EDTA, 0.5% IGEPAL CA-630 (Merck), 0.04 U/µL RNasin Plus RNase Inhibitor (Promega)], and the supernatants were used for binding to the beads. The beads were washed five times with wash buffer [10 mM Tris-HCl (pH 7.5), 150 mM NaCl, 0.5 mM EDTA], followed by elution as described above.

#### Cross-linking immunoprecipitation

Cross-linking immunoprecipitation was performed as described previously with some modifications^78^. OSCs had their medium removed and were UV cross-linked with 200 mJ/cm^2^ of 254 nm UV light on ice. Cells were then resuspended in CLIP-lysis buffer [20 mM HEPES-KOH (pH 7.3), 150 mM NaCl, 1 mM EDTA, 1 mM DTT, 0.5% NP-40, 2 μg/mL leupeptin, 2 μg/mL pepstatin A, 0.5% aprotinin] and treated with 10 U/μl RNase T1 (Thermo Fisher Scientific) for 15 minutes at room temperature. The supernatants were incubated with anti-Armi antibodies^20^ bound to Dynabeads Protein G (Thermo Fisher Scientific) at 4°C for 2 hours. Beads were washed three times with CLIP-wash buffer [20 mM HEPES-KOH (pH 7.3), 300 mM NaCl, 1 mM DTT, 0.05% NP-40, 2 μg/mL leupeptin, 2 μg/mL pepstatin A, 0.5% aprotinin], followed by three washes with high-salt wash buffer [20 mM HEPES-KOH (pH 7.3), 500 mM NaCl, 1 mM DTT, 0.05% NP-40, 2 μg/mL leupeptin, 2 μg/mL pepstatin A, 0.5% aprotinin]. The 5′ ends of the RNAs were radiolabeled with [γ-^32^P]-ATP (Revvity Health Sciences) in 1× PNK buffer using T4 polynucleotide kinase (New England Biolabs), followed by five washes with 1× PNK buffer without DTT. The Armi–RNA complexes were eluted with NuPAGE LDS Sample buffer (Thermo Fisher Scientific) and separated using NuPAGE Bis-Tris Gels (Thermo Fisher Scientific). The gels were stained with CBB, and the autoradiographs were detected using Typhoon FLA 9500 (Cytiva).

#### Prediction of Shu protein properties

The domain structure of Shu was predicted using HHpred^69^ with pfam30 as the HMM database and PsiBLAST as the MSA generation method. Domains longer than 10 amino acids with E-values less than 0.001 are shown^30^. The charge distribution was plotted using EMBOSS (window length 5)^79^. The potential of mean force over relative position vectors between two Shu molecules was predicted using FMAPB2 (pH 7.4; ionic strength 150 mM; 25°C)^44^, with the predicted structure obtained from the AlphaFold Protein Structure Database (AF-Q9W1I9-F1-model_v4)^68^ as input. The input structure of the ΔN77 mutant was generated by deleting the N-terminal 77 amino acids from the WT structure using CHARMM-GUI PDB Manipulator^80^. Interaction between intrinsically disordered regions were predicted using FINCHES^42^ with the Mpipi-GG forcefield^81^. Disorder predictions were also performed by FINCHES using metapredict V2-FF^43^.

#### Comparison of MOV10L1 homologs

Multiple sequence alignment was performed with Clustal Omega^70^ and visualized with ESPript 3.0^82^, employing the predicted structure of human MOV10L1 obtained from the AlphaFold Protein Structure Database (AF-Q9BXT6-F1-model_v4)^68^ for secondary structure depiction. Structure superposition was conducted and visualized with CueMol2 using secondary structure matching as the algorithm. The predicted structures of *Drosophila* Armi (AF-Q6J5K9-F1-model_v4), human MOV10L1 (AF-Q9BXT6-F1-model_v4), and human MOV10 (AF-Q9HCE1-F1-model_v4) were obtained from the AlphaFold Protein Structure Database^68^.

#### Protein purification

FLAG-tagged Armi WT and S769I mutant proteins were purified from S2 cells^83^ following previously established protocol^26^. Briefly, cells were lysed in Armi-lysis buffer [50 mM Tris-HCl (pH 8.0), 150 mM NaCl, 1 mM DTT, 0.1% NP-40, 1× cOmplete ULTRA Tablets (Merck), 25 µg/mL RNase A (Merck)], and the supernatant was incubated with Anti-FLAG M2 Affinity Gel (Merck), followed by washing with Armi-wash buffer [50 mM Tris-HCl (pH 8.0), 1 M NaCl, 1 mM DTT, 0.1% NP-40]. Armi was eluted by incubating the beads with 3 × FLAG peptide (Protein Ark) in Armi-elution buffer [50 mM Tris-HCl (pH 8.0), 150 mM NaCl, 1 mM DTT, 0.1% NP-40, 10% glycerol], after equilibrating the beads with Armi-elution buffer.

GST-tagged Shu proteins were expressed in *E. coli* [Rosetta 2(DE3) strain (Novagen)], and cells were lysed via sonication in Shu-lysis buffer [50 mM Tris-HCl (pH 8.0), 500 mM NaCl, 5 mM DTT, 1% Triton X-100] supplemented with 1× cOmplete ULTRA Tablets (Merck). Shu-GST in the supernatant was bound to Glutathione Sepharose 4B (Cytiva), washed with Shu-lysis buffer, and then with PBS. For pull-down assays, the washed beads were used directly as bait without elution. For *in vitro* dynamics assays, Shu was cleaved from the GST tag and eluted by incubating the beads with GST-HRV3C (purified from *E. coli* in-house) at 4°C overnight. The eluted proteins were further purified using an ENrich SEC 650 column (Bio-Rad) with the NGC Quest 10 Plus Chromatography System (Bio-Rad).

#### In vitro dynamics assay

Proteins were labeled with ATTO550 maleimide (ATTO-TEC), quenched with excess DTT, and the free dyes were removed using Zeba Spin Desalting Columns (Thermo Fisher Scientific). Proteins diluted in PBS were mixed with free ATTO488 maleimide pre-quenched with DTT to correct for changes in viscosity^84^. Distribution coefficients were collected using an LSM 980 laser scanning confocal microscope with Dynamics Profiler (Carl Zeiss).

#### Pull-down assay

Recombinant Armi proteins were incubated at 4°C for 2 hours with Shu-GST bound to Glutathione Sepharose 4B in Armi-elution buffer (see above) supplemented with 0.17 µg/µL total RNA extracted from OSCs. The beads were washed five times with Armi-elution buffer. Proteins were eluted in 2x Laemmli SDS sample buffer and analyzed by manual western blotting (see above).

#### RNA unwinding assay

The RNA unwinding assay was performed as previously described with some modifications^26^. Synthetic RNA oligonucleotides (Merck) were purified by denaturing PAGE, and the shorter oligo was radiolabeled with [γ-^32^P]-ATP in 1×PNK buffer using T4 polynucleotide kinase. The labeled oligo was mixed with an unlabeled longer oligo and annealed in annealing buffer [20 mM Tris-HCl (pH 7.5), 50 mM NaCl, 1 mM EDTA]. 50 nM of recombinant Armi proteins were bound with 1 nM of RNA duplexes in unwinding buffer [50 mM Tris-HCl (pH 7.5), 50 mM KOAc, 1 mM DTT, 0.01% NP-40, 1 U/μL RNasin Plus RNase Inhibitor] for 5 minutes at 25°C. The unwinding assay was initiated by adding MgCl_2_, ATP, and an unlabeled shorter oligo to a final concentration of 0.5 mM. The reaction was stopped by adding the same volume of quench buffer [50 mM EDTA, 0.5% SDS, 0.04% bromophenol blue, 0.04% xylene cyanol, 20% glycerol], and the mixture was separated by non-denaturing PAGE. Autoradiographs were detected using Typhoon FLA 9500. The sequences of the oligos used are shown in Table S1.

#### Quantification and statistical analysis

For manual western blotting, signal intensities were quantified using ImageJ^71^. Relative pull-down/co-immunoprecipitation (co-IP) efficiencies were calculated by dividing the signal intensity of the prey bound to beads by that of the bait on beads, and then by the intensity of the prey in the input. In immunofluorescence experiments, cells were manually counted. Sample sizes are described in figure legends, and data are presented as the mean ± SEM. For the *in vitro* dynamics assay, R^72^ with MASS library^73^ was used for linear regression to calculate *k_D_* values with standard errors. Comparisons of *k_D_* values were performed using Welch’s t-test in Microsoft Excel (Microsoft). Relative pull-down/co-IP efficiencies were compared using the Bootstrap method with BootstRatio^74^. p values less than 0.05 were considered as statistically significant.

---

**Table S1.**
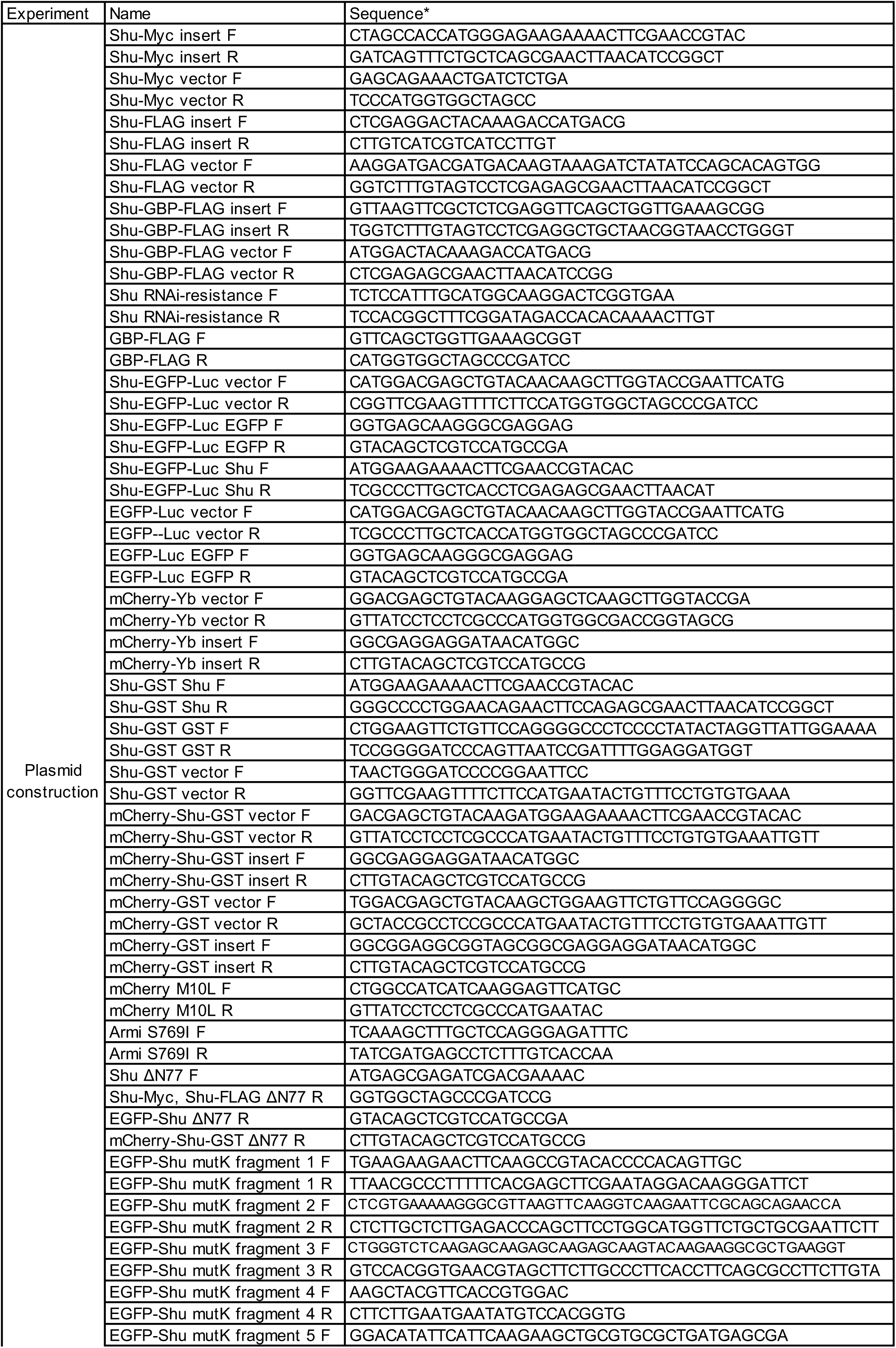
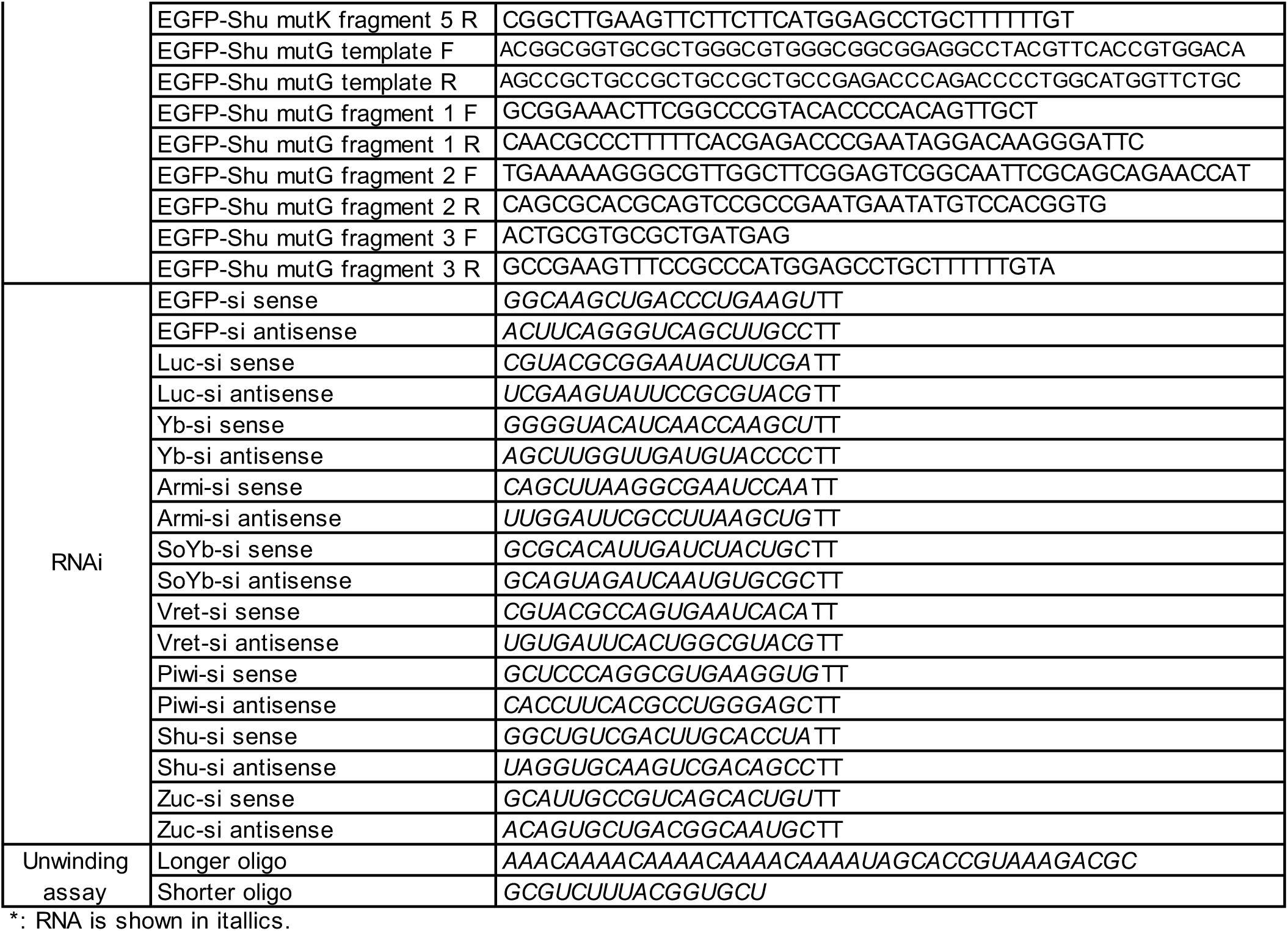
Sequences of oligonucleotides.

